# Regulatory sites of CaM-sensitive adenylyl cyclase AC8 revealed by cryo-EM and structural proteomics

**DOI:** 10.1101/2023.03.03.531047

**Authors:** Basavraj Khanppnavar, Dina Schuster, Pia Lavriha, Federico Uliana, Merve Özel, Ved Mehta, Alexander Leitner, Paola Picotti, Volodymyr M. Korkhov

**Affiliations:** Laboratory of Biomolecular Research, Division of Biology and Chemistry, Paul Scherrer Institute, Villigen, Switzerland; Department of Biology, Institute of Molecular Biology and Biophysics, ETH Zurich, Switzerland; Department of Biology, Institute of Molecular Systems Biology, ETH Zurich, Switzerland; Department of Biology, Institute of Biological Chemistry, ETH Zurich, Switzerland

## Abstract

Membrane adenylyl cyclase AC8 is regulated by G proteins and calmodulin (CaM), mediating the crosstalk between the cAMP pathway and Ca^2+^ signalling. Despite the importance of AC8 in physiology, including cognitive functions and memory, the structural basis of its regulation by G proteins and CaM is not well defined. Here we report the 3.5 Å resolution cryo-EM structure of the bovine AC8 bound to Ca^2+^/CaM and the stimulatory Gαs protein. The structure reveals the architecture of the ordered AC8 domains bound to Gαs and a small molecule activator forskolin. The extracellular surface of AC8 features a negatively charged pocket, a potential site for unknown interactors. Despite the well resolved forskolin density, the captured state of AC8 does not favour tight nucleotide binding. The structural proteomics approaches, limited proteolysis and crosslinking mass spectrometry, allow us to identify the contact sites between AC8 and its regulators, CaM, Gαs, and Gβγ, as well as to infer the conformational changes induced by these interactions. Our results provide a framework for understanding the role of flexible regions in the mechanism of AC regulation.

## INTRODUCTION

Membrane-integral adenylyl cyclases (ACs) catalyse the conversion of ATP into a universal second messenger, cyclic AMP (cAMP), and play a significant role in transmitting information from sensory cell surface receptors, including G protein-coupled receptors (GPCRs), to multiple effector proteins such as protein kinase A or cyclic nucleotide-gated ion channels (*1, 2*). Membrane ACs (AC1-9) share a common structural blueprint: each is a polytopic membrane protein with twelve transmembrane helices and two catalytic domains, C1 and C2 (*2*). They show ∼50 % homology (*3*). The modes of regulation and tissue expression vary for each AC type (*4*). All isoforms are activated by the stimulatory G protein alpha (Gαs) subunit (*5*), and some isoforms (AC1, AC5 & AC6) are inhibited by the inhibitory G protein alpha (Gαi) subunit (*6*). Gβγ subunits have been shown to modulate the cyclase activity in an AC isoform-specific manner, with some isoforms (AC2, AC4, AC5, AC6, AC7) being activated (*7-10*) and some isoforms (AC1, AC3, AC8) being inhibited by Gβγ (*7, 11-13*), with AC9 apparently not sensitive to Gβγ (*7*).

The molecular basis of the conversion of ATP to cAMP by membrane ACs was revealed by X-ray crystallographic studies of the chimeric catalytic domains of AC5_C1_/AC2_C2_ (*14-16*). These studies defined the structural basis of how ACs perform a two-metal catalysis reaction, how they are activated by a Gαs subunit and how the plant-derived small molecule forskolin activates the enzyme. Forskolin and Gαs can activate membrane ACs independently or synergistically through stabilising the interaction between the catalytic domains C1a and C2a (*15*). The mechanism of regulation by other G protein subunits, such as Gβγ, is not fully understood, partly due to the lack of structural data. Peptide-based inhibition studies have suggested that Gβγ binds the C1/C2 catalytic domains of AC2 (*17*), AC5 (*18*) and AC6 (*19*). Additionally, Gβγ has been reported to interact with the N-termini and/or the C1b domain of different AC isoforms, including AC1 (*18*), AC2 (*13*), AC5 (*18, 19*), AC6 (*10, 19*).

The Ca^2+^/CaM-sensitive ACs, AC1 (*20*), AC3 (*21*), and AC8 (*20*) play a pivotal role in cellular signalling (*1, 22*). They mediate the crosstalk between distinct signalling pathways (i.e. the GPCR/cAMP pathway and the Ca^2+^ signalling pathway), integrating the different signalling inputs and fine-tuning the cellular responses to them (*22*). The effect of Ca^2+^ on ACs is mediated by the Ca^2+^-sensor protein CaM (*23, 24*). Binding of CaM to Ca^2+^/CaM-sensitive ACs stimulates their catalytic activity in a Ca^2+^-dependent manner. AC8 was shown to interact with CaM via two distinct conserved calmodulin binding domains: an amphipathic CaM-binding motif at the N-terminus and an IQ-like CaM-binding motif in the cyclase C2b domain (**Fig. 1a**) (*25, 26*). Based on *in vitro* studies, the N-terminal CaM-binding motif was proposed to contribute to the constitutive tethering of CaM to AC8, while not being absolutely required for stimulation of the catalytic activity (*27-29*). The CaM-binding motif on the AC8 C2b domain, on the other hand, was suggested to be critical for activation of AC8 by Ca^2+^/CaM (*25, 26, 30*). Similar to AC9, the deletion of the C2b domain results in constitutive superactivation of AC8, and it has been proposed that the Ca^2+^/CaM complex may be involved in the regulation of the autoinhibitory mechanism of AC8 (*25, 26, 30*).

**Figure 1.**
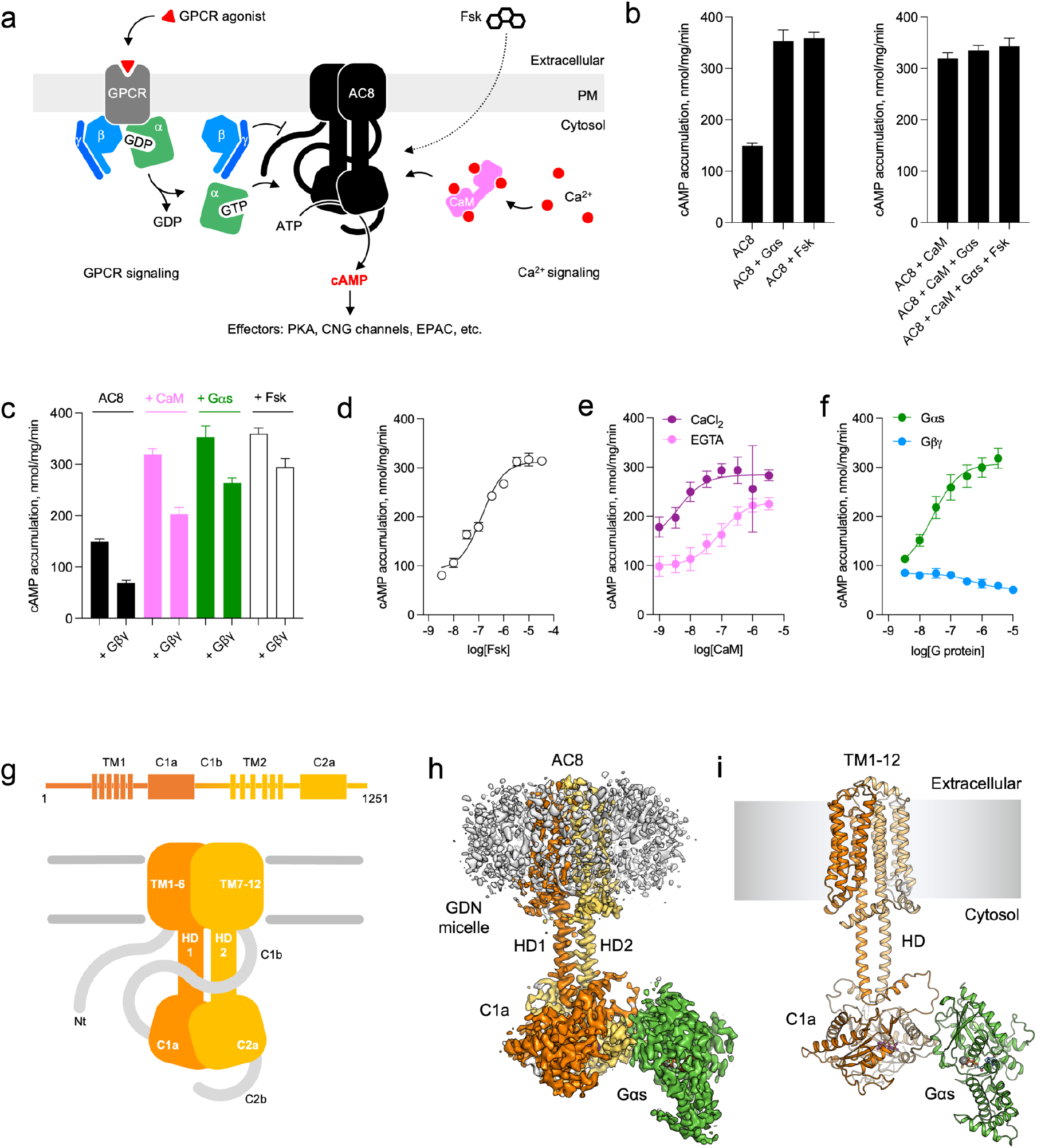
Biochemical characterisation of purified bovine AC8. **(a)** Schematic diagram depicting the key players in regulation of AC8 and its role in mediating the crosstalk between the cAMP and Ca^2+^ signalling pathways; Fsk – forskolin, CaM – calmodulin, PM – plasma membrane. **(b-c)** Enzymatic activity of AC8 in the presence of various interacting partners (CaM, Gαs, Gβγ and forskolin); n = 4. The AC8 control bar is identical in b and c. **(d-f)** Dose-response curves for AC8 activity in the presence of various interaction partners. For all experiments, the data are shown as mean ± S.D. (n = 3). **(g)** Pictographic representation of membrane adenylyl cyclase AC8 topology. **(h-i)** Cryo-EM map **(h)** and model **(i)** of the AC8-Gαs-Ca^2+^/CaM-Forskolin-MANT-GTP complex.

We have previously determined the high-resolution structure of the full-length bovine AC9, revealing molecular details of AC9 autoinhibition by its C-terminal domain C2b (*31*). Despite the availability of a homologous AC9 structure and existing biochemical data on AC8, our understanding of the molecular mechanisms of Ca^2+^/CaM-sensitive AC regulation by CaM and G protein subunits is incomplete. To address this knowledge gap, we determined the structure of purified bovine Ca^2+^/CaM-sensitive AC8, complemented by mass spectrometry-based structural proteomics approaches (XL-MS and LiP-MS) and functional studies. The combination of these techniques allowed us to gain deeper insights into the regulation of AC8 and better understand the role of its structured and flexible domains.

## RESULTS

### Purification and biochemical characterisation of bovine AC8

AC8 is known to interact with and to be regulated by CaM, Gαs, Gβγ and forskolin (**Fig. 1a**) (*1, 7, 20, 25, 30*). To gain insight into the structure, function and regulation of AC8 (**Fig. 1a**), we expressed bovine AC8 in the human embryonic kidney 293 (HEK293) GnTI^-^ cells using a tetracycline-inducible expression system, purified the protein in detergent by affinity chromatography (**Fig. S1**), and assessed functional integrity of the preparation with AC activity assays **(Fig. 1b-c)**. Quantitative mass-spectrometry (MS) -based assessment of the purified AC8 preparations determined that endogenously expressed CaM is a negligible contaminant in our AC8 preparations (AC8 : CaM ratio of ∼500 : 1; **Fig. S2**).

The purified AC8 was efficiently activated by its endogenous activators CaM and Gαs, as well as by the small molecule activator forskolin (**Fig. 1d-f**). The EC_50_ of CaM-activation was 4.1 nM and 86.6 nM in the presence and absence of CaCl_2_, respectively (**Fig. 1e**). Thus, our *in vitro* experiments using full-length AC8 confirm that CaM-dependent activation of AC8 is a Ca^2+^-dependent process, as has been observed previously (*20, 25, 30*). Fluorescence-detection size-exclusion chromatography (FSEC) binding assays using purified AC8 and a C-terminally yellow fluorescent protein (YFP) tagged CaM (CaM-YFP) showed that the apparent affinity of CaM-YFP for AC8 is ∼70 nM in the presence of Ca^2+^ (**Fig. S3**). In the presence of a Ca^2+^ chelating agent, EGTA, high affinity binding could not be observed in the FSEC experiment. The difference between the AC8 activation EC_50_ values and the apparent K_d_ values in the binding assays likely stems from the use of a fluorescently tagged CaM and the non-equilibrium separation technique (FSEC). Nevertheless, the adenylyl cyclase activity assays combined with the binding assays clearly indicate that in the presence of Ca^2+^ CaM binds AC8 with nanomolar affinity.

The GTPγS-bound Gαs activated AC8 with high affinity (EC_50_ of 25.5 nM; **Fig. 1f**). AC8 was inhibited by Gβ_1_γ_2_ subunits, consistent with previous observations (*7*). Interestingly, the EC_50_ for the activation of AC8 by Gαs is 30-fold lower than for bovine AC9, suggesting an isoform-dependent affinity. In contrast to CaM and Gαs, Gβγ inhibits AC8 with lower micromolar affinity with an IC_50_ of 0.4 μM (**Fig. 1e-f**). Forskolin activates AC8 with nanomolar affinity (EC_50_ = 136 nM) (**Fig. 1d**). The results are in line with the available body of evidence (*1, 7, 20, 25, 30*), confirming that our purified AC8 preparations represent a native like state of the protein, with intact sensitivity to the known regulators.

### Cryo-EM structure of the AC8-CaM-Gαs complex

To structurally characterise AC8, we used single-particle cryo-EM analysis of the protein in the presence and in the absence of Ca^2+^/CaM, Gβγ and Gαs. The concentrations of each of the protein components (AC8, CaM, Gαs) in the cryo-EM samples were controlled by mixing the purified proteins at specific molar ratios immediately prior to grid freezing. Among the tested combinations, one sample proved to be suitable for structure determination: AC8 in a complex with Ca^2+^/CaM and activated Gαs, in the presence of MANT-GTP and a forskolin. After multiple rounds of 2D and 3D classifications, the best set of particles resulted in a 3D reconstruction at a resolution of 3.5 Å (**Fig. S4, S5 & S6, Table S2**).

The structure of AC8-CaM-Gαs revealed most of the features of AC8, including the transmembrane (TM) domain, the helical domain (HD), the catalytic domain (CAT), as well as the G protein Gαs subunit (**Fig. 1g-i**). The N-terminus (1-164), the C-terminus (1165-1251) and the C1b domain (585-670) could not be resolved in the final 3D reconstruction.

### AC8-Gαs interface

The stimulatory G protein subunit Gαs activates each of the membrane AC isoforms. We compared the features of the AC8-Gαs interaction revealed by our structure with previously reported structures: the cryo-EM structure of AC9-Gαs (*31, 32*) and the X-ray crystallography structure of the AC2_(C2a)_-AC5_(C1a)_-Gαs complex (*14*). Each of these ACs interacts with Gαs via conserved and non-conserved polar residues (**Fig. S10, Table S3**). However, we can observe differences in the number and type of interfacial residues (**Fig. S10**). Although these residues are not necessarily involved in direct interactions with Gαs (except for a few hydrophobic contacts), they may influence the AC-G protein binding. Interestingly, the apparent affinity of Gαs for AC8 (EC_50_ of 25.5 nM, **Fig. 1f)** is higher than that for AC9 (EC_50_ of 802.9 nM) (*32*). The difference in affinity between Gαs and different AC isoforms is likely explained by the divergent interfacial interactions, which may play a role in a complex cellular environment where specific signalling inputs may be required, e.g., in the presence of several co-expressed AC isoforms.

### Conformational heterogeneity in the substrate binding site of AC8

The structure showed a clear density corresponding to forskolin (**Fig. 2a-c, Fig. S5**). In stark contrast, the density for MANT-GTP was not well-resolved (**Fig. 2a-c, Fig. S5**). We tested whether this could be explained by a drop in affinity for MANT-GTP in the presence of forskolin, but the affinity of AC8 for MANT-GTP was not affected by the presence of either forskolin or other activators (**Fig. 2d-e**). We previously observed that AC9 is capable of binding MANT-GTP in different conformations dependent on the presence of forskolin (*32*). These observations are in contrast to the distinct single conformation of MANT-GTP observed in the crystal structure of the chimeric catalytic domain of AC2_(C2a)_-AC5_(C1a)_ (*14*). Thus, the behaviour of the nucleotide binding site in AC8 is distinct from that of AC9 or the chimeric AC2_(C2a)_-AC5_(C1a)_.

**Figure 2.**
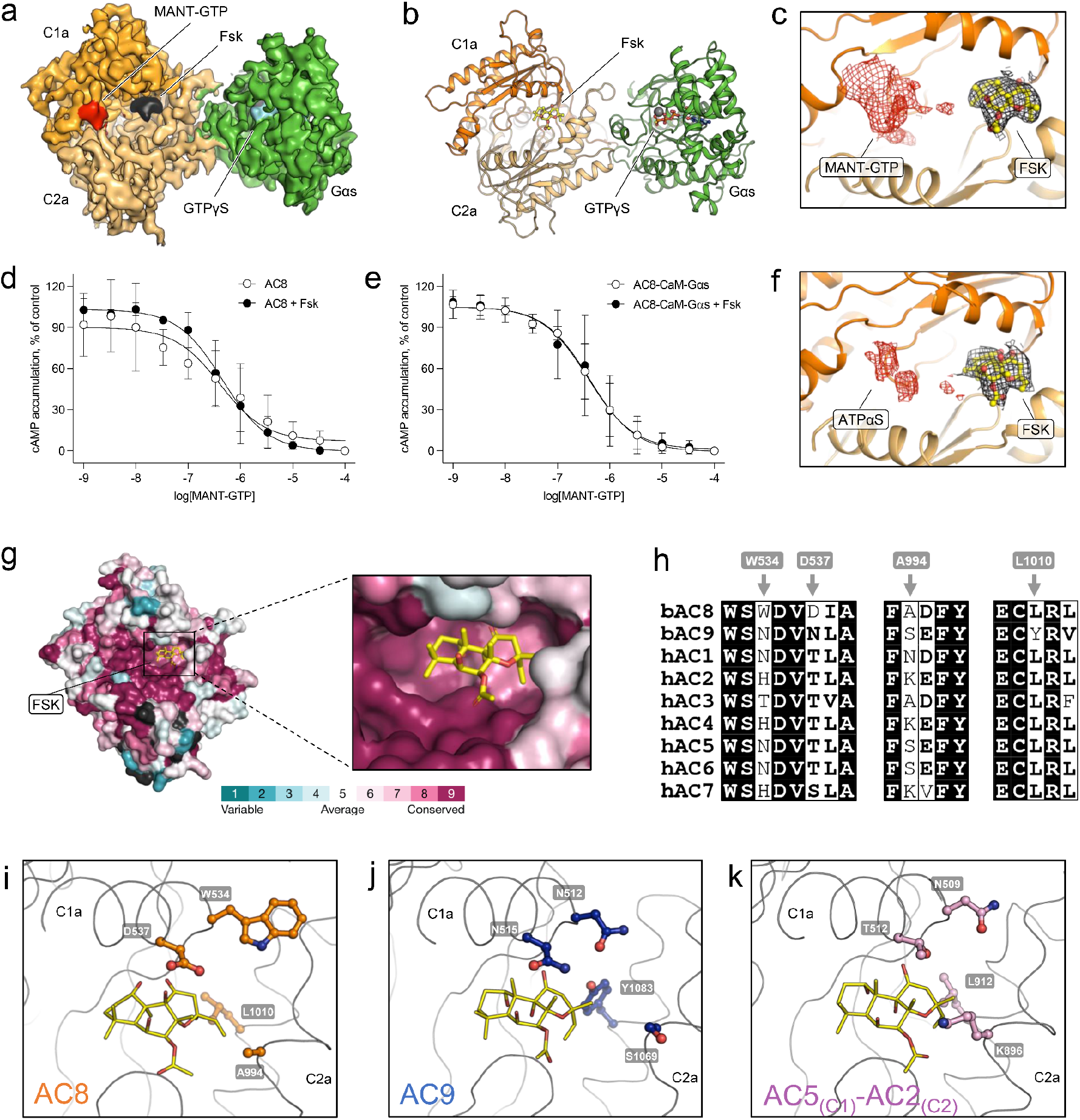
ATP- and forskolin-binding pockets in AC8. **(a, b)** Cryo-EM map and model of the catalytic domain of AC8 in complex with Gαs, Ca^2+^/CaM, forskolin and MANT-GTP. **(c)** Electron density in the substrate-binding pocket of AC8 showing a well-defined density for forskolin and a poorly resolved density for MANT-GTP. **(d-e)** Dose-response curves of MANT-GTP in the presence and absence of forskolin, CaM, and Gαs. For all experiments, the data are shown as mean ± S.D. (n = 3). **(f)** Unresolved density in the ATP-binding site of the AC8-Gαs-Ca^2+^/CaM-forskolin-ATPαS complex solved in lipid nanodiscs. **(g)** Conservation scores for nine membrane AC isoforms computed with ConSurf (*46*). **(h)** A segment of multiple sequence alignment showing key non-conserved residues in forskolin binding interfaces in nine membrane AC isoforms. **(i-k)** Comparison of forskolin-binding interfaces in AC8, AC9 (PDB ID: 6r40), and the chimeric AC2_(C2a)_-AC5_(C1a)_ complex (PDB ID: 1tl7). The interfacial residues in panels h-k were chosen based on their proximity (within 4Å) to forskolin in any one of the AC isoforms.

To ensure that the observed conformation is not induced by the detergent micelle, we reconstituted AC8 in MSP1E3D1 nanodiscs with native brain polar lipids and prepared a cryo-EM sample using similar conditions **(Fig. S7a-b)**. As the density of MANT-GTP was not well resolved in the detergent-purified structure, we replaced MANT-GTP with ATPαS. We reasoned that ATPαS should be a better substrate surrogate than the nucleotide-based inhibitor MANT-GTP since it is structurally similar to the substrate ATP. The lipid nanodisc sample was processed similar to the detergent sample. After multiple rounds of 2D and 3D classifications, the 3D reconstruction resulted in a 3.97 Å map of the AC8-Gαs complex and a 3.5 Å resolution structure after focused refinement of the AC8 catalytic domain-Gαs complex. The conformation of the nanodisc reconstituted AC8 is similar to the conformation observed in detergent. The N-terminus, C-terminus, C1b, and CaM remain unresolved **(Fig. S7, S8 & S7**). The density for ATPαS was also not well-resolved in this structure **(Fig. 2f & Fig. S9)**. Together with the lack of a well-resolved density of MANT-GTP, these data are indicative of substrate and inhibitor conformational heterogeneity.

### Specificity of forskolin binding to AC8 and other membrane ACs

The forskolin-binding interfaces of AC8, AC9 and AC2_(C2a)_-AC5_(C1a)_ show both conserved and non-conserved interfacial residues **(Fig. 2g-k)**. The interfacial residues closer to the ATP-binding site are highly conserved, whereas the residues near the Gαs binding site exhibit some variability **(Fig. 2g)**. These minor differences may contribute to the observed differences in affinity of forskolin for the various AC isoforms (*33, 34*), and may also underlie the Gαs-dependent conditional activation of AC9 (*31, 35*). Comparisons of the forskolin binding sites of AC8, AC9 and AC2_(C2a)_-AC5_(C1a)_ show that functional differences between these ACs are likely to be explained by four residues (in AC8: W534, D537, A994, and L1010) **(Fig 2h-k)**. All four are in close proximity of forskolin in one or more of the available AC structures. One residue in particular, L1010, stands out: it is conserved among all ACs but one, AC9. In AC9 this position is occupied by Y1083, which directly faces forskolin with its hydroxyl group (**Fig. 2h-k**). AC9 alone is not stimulated by forskolin and can only be activated by forskolin in the presence of a bound Gαs subunit (*32, 35*). It is thus likely that a leucine residue in this key position (L1010 in AC8) favours G protein-independent forskolin activation of a membrane AC, whereas the presence of a tyrosine residue (Y1083 in AC9) may require G protein to enable forskolin-mediated AC activation.

### TM domain of AC8 features a putative pocket for extracellular ligands

Comparison of the structure of AC8 with the previously solved full-length structures of the membrane ACs, AC9 and the mycobacterial membrane AC Rv1625c/Cya, showed differences in the degree of rotation between the CAT and TM domains (**Fig. 3a**). Nevertheless, the AC8 TM domain adopts a similar 12 TM helix arrangement to that of AC9, and a similar organisation of TM1-5/TM7-11 to TM1-5 of Cya (**Fig. 3b-d**). The observed differences in the relative orientation of the TM domain and CAT with respect to each other, after 3D alignment of AC8 and its homologues, seems to originate from the coiled-coiled HD helices of the ACs (**Fig. 3a**). The HDs of AC8 and AC9 are substantially longer than HD of Cya, and the difference in orientation of the TM domains relative to the CAT domains in AC8 and AC9 may be due to small structural changes in the HDs.

**Figure 3.**
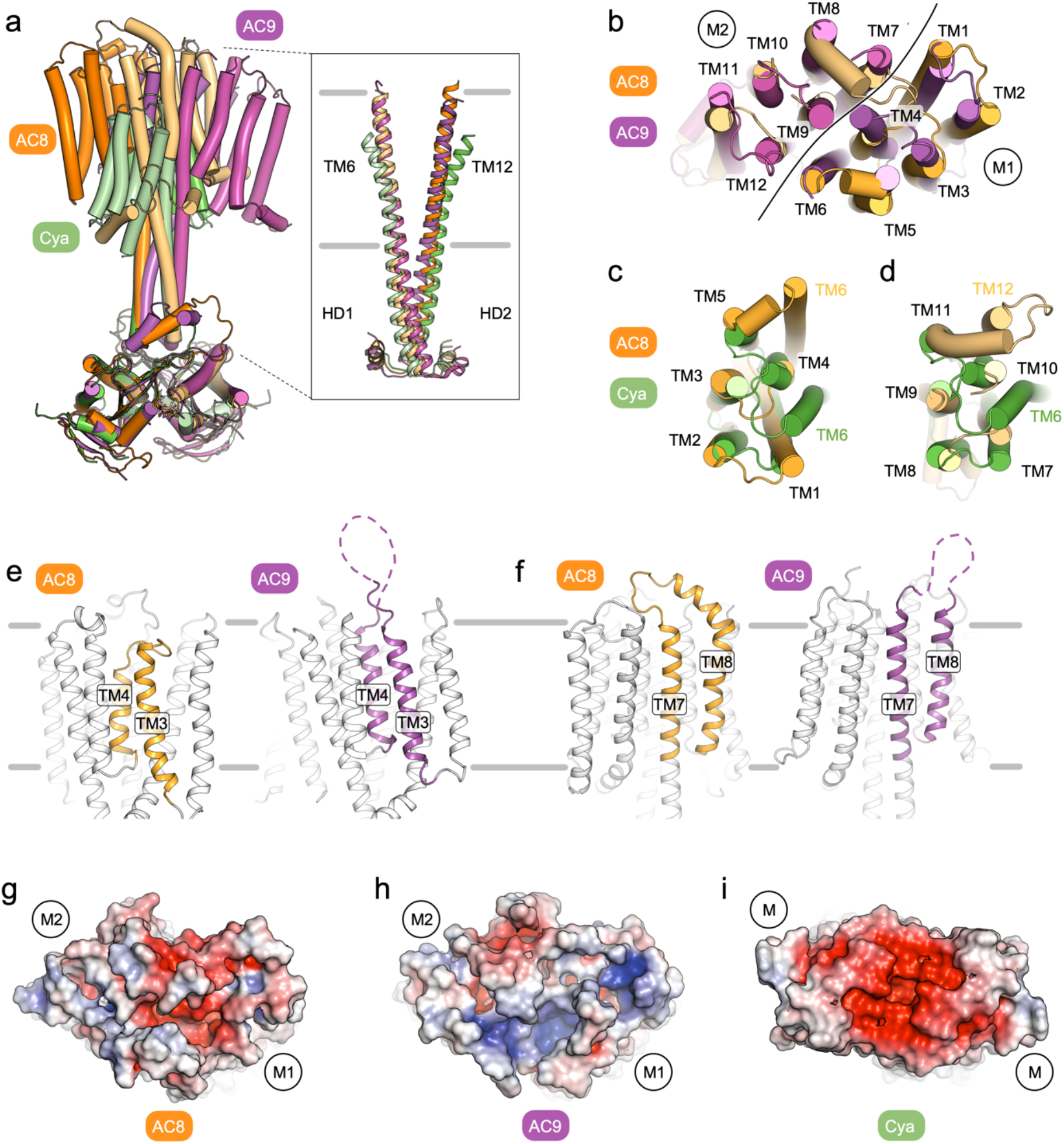
Key structural features of the transmembrane domain of AC8. **(a)** Comparison of cryo-EM structures of AC8, AC9 and *M. tuberculosis* Rv1625c/Cya. The membrane domains of these ACs are tilted differently relative to the helical and catalytic domains. The insert shows the alignment of the helical domains of the three membrane ACs of the known structure. **(b-d)** Alignment of the membrane domains of AC8 and AC9 (b) and AC8 and Cya (c-d). The line in b indicates the boundaries of the two hexahelical domains (“M1” and “M2”). **(e-f)** Depiction of the key structural differences at the extracellular side of the AC8 TM domain compared to AC9, in TM3-4 (short loop in AC8, disordered loop in AC9) and in TM7-8 (ordered folded loop in AC8, disordered extended loop in AC9). **(g-i)** An extracellular view of the TM domains of AC8, AC9, and Rv1625c/Cya showing the potential extracellular ligand binding pockets in surface representation, coloured according to electrostatic potential. The “M1” and “M2” labels indicate the positions of the 6-TM bundles of AC8 (g) and AC9 (h), “M” denotes the 6-TM bundles of Rv1625c/Cya (i).

The extracellular region (except for residues 811-831) of the AC8 TM domain is better resolved compared to that in AC9 (**Fig. 3e-f, Fig. S5**). This near-complete structure of the extracellular side of the TM domain reveals a prominent cavity formed at the interface of the TM1-7 helices (**Fig. 3g**). The pocket is formed by the shorter TM4 helix and loop between TM3-4 (ECL2, **Fig. 3e-h, Fig. S5**) and is partially negatively charged, reminiscent of Cya (**Fig. 3i**) (*36*). Our previous study of Rv1625c/Cya (*36*) indicated that the membrane anchor of this mycobacterial membrane AC may have a receptor function for cations, such as divalent metals. The features of the extracellular portion of the TM region in AC8 are compatible with a similar function, i.e. interactions with positively charged ligands (ions, small molecules, peptides or proteins). Further studies will be required to determine whether the extracellular surface of AC8 may contain functionally relevant ligand-binding sites.

### LiP-MS reveals involvement of flexible domains in activating and inhibiting interactions of AC8

Limited proteolysis-coupled mass spectrometry (LiP-MS) has previously been demonstrated to detect protein-binding interfaces upon addition of a protein interactor of interest to a complex proteome or membrane extract, due to changes in protease accessibility upon protein-protein interaction (*37*). We therefore used LiP-MS to detect binding interfaces and other regions of altered surface accessibility upon interaction of AC8 with Gαs and Gβγ. We also reanalysed a previously published LiP-MS dataset of the interaction of AC8 with CaM, conducted under the same experimental conditions (PRIDE ID: PXD039520; (*37*)). Since binding interfaces of AC8-CaM were previously known, and since we determined those of AC8-Gαs experimentally by cryo-EM, we could use these orthogonal datasets for cross-validation.

We prepared crude membrane suspensions of AC8-YFP-overexpressing HEK 293 GnTI-cells and titrated them with interactors of interest in increasing concentrations. We performed limited proteolysis with proteinase K on all samples under native conditions, followed by denaturation and complete trypsin digestion. We controlled for protein abundance changes control samples subjected only to the trypsinisation step under denaturing conditions after treatment with the highest or lowest interactor concentrations (see Materials and Methods). We measured all samples with data-independent acquisition and fit four-parameter dose-response curves onto the measured peptide profiles. To determine regions changing in protease accessibility, we filtered for peptides with a Pearson correlation r of >0.85 between the measured peptide intensities and the fitted dose-response model (**Fig. S11**). Peptides with a correlation r <0.85 were considered unchanged (**Fig. 4d**). This threshold (0.85) was defined based on the validation AC8-CaM and AC8-Gαs data sets, for which the binding interfaces were known. We detected 167 AC8-YFP peptides in the CaM dataset, 177 in the Gβγ dataset, and 180 in the Gαs dataset, corresponding to a sequence coverage of 55-60%. As expected, transmembrane domains of AC8 were not detected in any of the datasets. Nine AC8 peptides showed evidence of a change in protease accessibility upon addition of CaM (**Fig. 4a**), six peptides upon addition of Gαs (**Fig. 4b**), and six different peptides upon addition of Gβγ (**Fig. 4c**).

**Figure 4.**
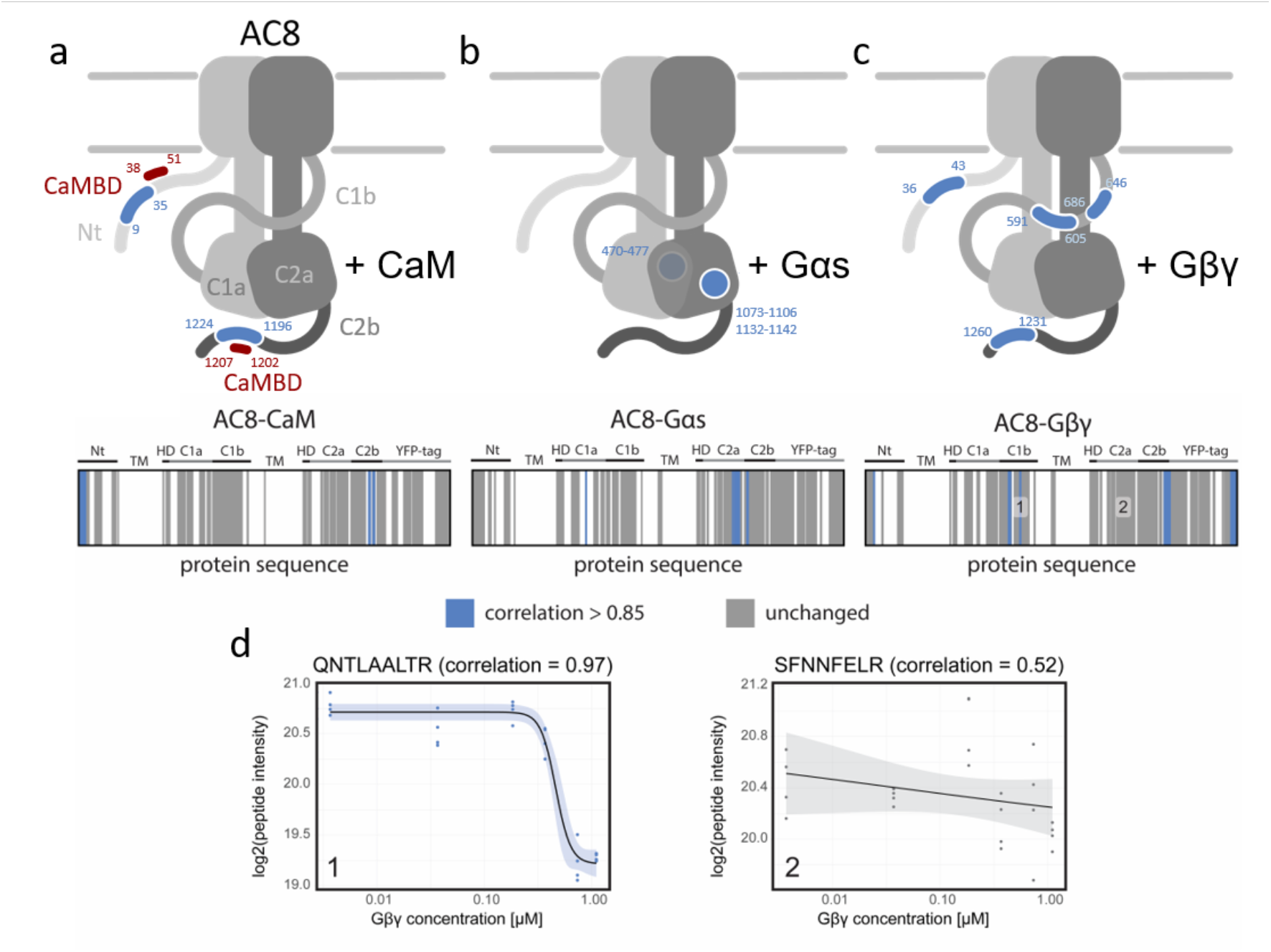
LiP-MS reveals binding interfaces of AC8 interactors. **(a-c)** Sketches of AC8 structures with detected changes highlighted with blue lines (on flexible regions) or circles (on structured regions). CaM-binding domains are indicated in red. Barcode plots in the middle of the figure show detected peptides in grey and peptides that changed upon addition of an interactor (Pearson correlation r > 0.85) in blue, along the sequence of the protein. **(d)** Two exemplary interaction curves for peptides detected in the AC8-Gβγ experiment are shown below. The significantly changing peptide is highlighted in blue, the unchanged peptide is shown in grey. Addition of CaM has an effect on protease accessibility of the AC8 N-terminus and the AC8 C-terminus. Addition of Gαs changes the accessibility of regions in the catalytic domains C1a and C2a, with one peptide overlapping between the C-terminal end of C2a and C2b. Gβγ addition changes the accessibility of regions in the three flexible protein domains: the N-terminus, C1b and C2b with additional changes on the YFP-tag.

The changes induced by the different interactors were located on different AC8 domains. CaM induced accessibility changes on the AC8 N-terminus and C2b domain (C-terminus), at the known binding interfaces (*29, 30*) (**Fig. 4a**). Interestingly, the significantly changing peptides show profiles with opposite directionality. We find that peptides closer to the CaM binding motifs increase in abundance, while peptides further away decrease in abundance. Assuming that the peptides at the binding sites increase in abundance due to increased protection from proteolytic cleavage, the regions further away likely become more accessible to proteolytic cleavage and therefore we detect a decrease in their abundance upon CaM binding. Addition of Gαs to AC8 induced changes in both catalytic domains C1a and C2a, with the one peptide at the C-terminal end of C2a, overlapping with the first amino acids of C2b. The detected accessibility changes cover the binding interface we could resolve in our cryo-EM structure on C2a (residues 1073-1081), as well as an extension of this region (1083-1106), and a neighbouring peptide (1132-1142) (**Fig. 4b**). We also identify an accessibility change on C1a (470-477) that could result from the Gαs N-terminus that is not resolved in our cryo-EM structure. Addition of Gβγ changed the surface accessibility of all three flexible AC8 domains (N-terminus, C1b and C2b domains) and the YFP-tag (**Fig. 4c**). The changes detected upon interaction with Gβγ are distinct from the changes induced by CaM or Gαs. Our data shows that CaM, Gαs and Gβγ interact with and/or induce conformational changes in distinct domains of AC8, and that these interactions involve flexible regions of the cyclase. Moreover, these results indicate that the AC8 activation mechanisms of CaM and Gαs are distinct from each other and may occur concurrently, in line with the activity data shown in **Fig. 1** and consistent with the role of AC8 as an integrator of the G protein and the Ca^2+^ signalling inputs.

### XL-MS provides evidence of AC8 oligomerisation

We used XL-MS to better interpret the changes detected with LiP-MS, in particular to distinguish conformational changes from binding interfaces and additionally to obtain proximity information. The results of the two crosslinkers we used (dissuccinimidyl suberate (DSS) (*38*) and a combination of pimelic acid (PDH) and the coupling reagent 4-(4,6-dimethoxy-1,3,5-triazin-2-yl) (DMTMM) (*39*)) are combined and discussed together.

We first examined crosslinks between AC8 subunits, since previous studies have suggested that ACs are able to form homodimers with interaction interfaces located in their catalytic and transmembrane domains (*40, 41*). Consistent with these results, our XL-MS experiments of AC8 identified several homomultimeric links (between two separate AC8 molecules), indicating self-association of AC8 (**Fig. 5b**). The general architecture and homomultimeric link sites are reproducible across the different screened conditions. These sites are found on the structured (C1a, C2a) domains, as well as the flexible domains (N-terminus, C1b and C2b domains). Docking of a dimeric structure based on the identified self-association links is not possible, indicating either a high flexibility of AC8 dimers or the existence of different dimeric or oligomeric structures. Consistent with these data, the purified AC8 elutes as two peaks (monomer and dimer) in our SEC chromatogram (**Fig. S1**). Future work on stabilising or isolating a conformationally homogeneous dimeric AC8 will be necessary to determine the structure of the dimer by cryo-EM. The XL-MS results presented here serve as an important piece of evidence for the existence of dimeric or multimeric AC assemblies in the absence of direct structural evidence.

**Figure 5.**
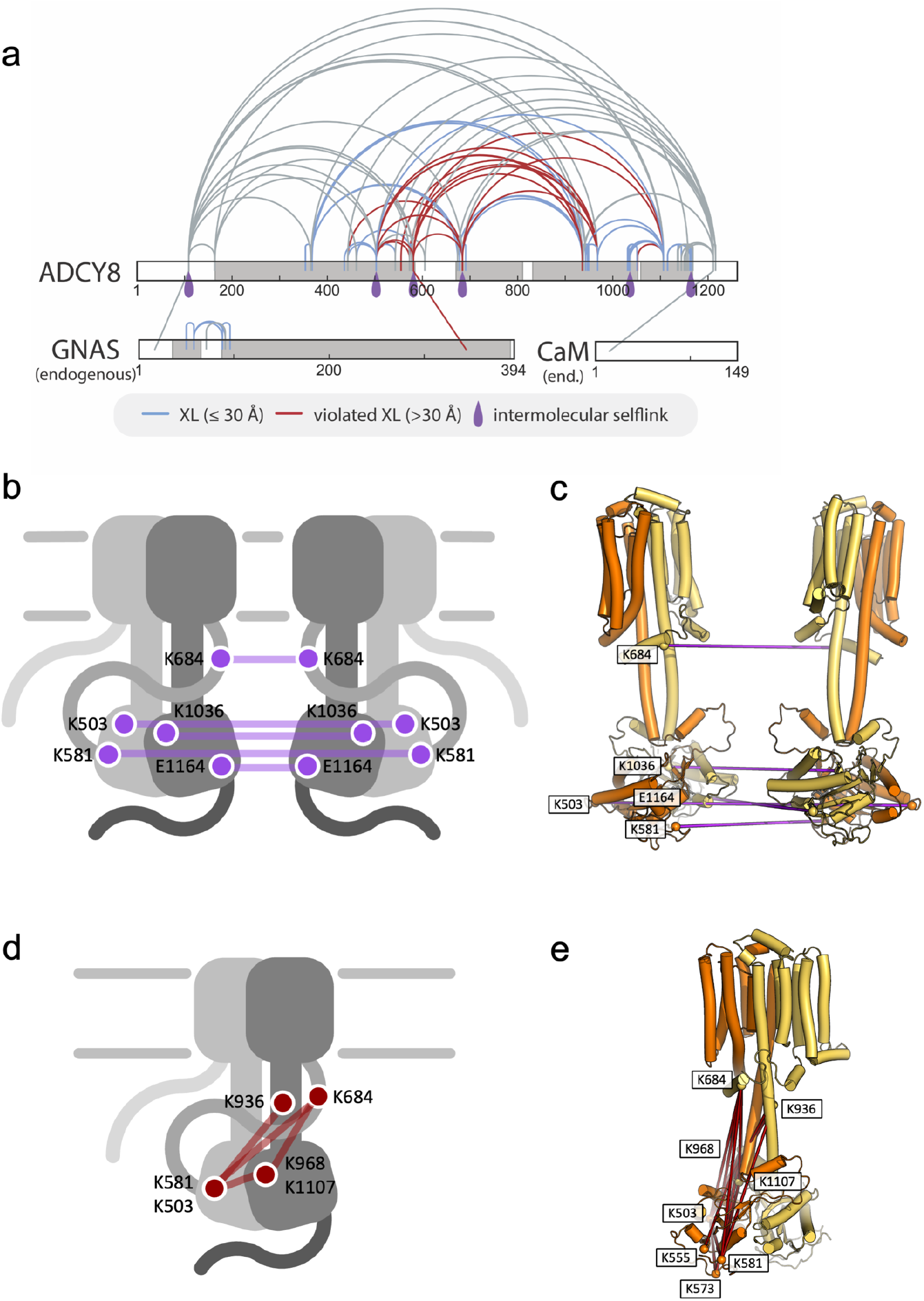
Intermolecular self-links and crosslinks between AC8 and copurified interactors indicate flexibility in AC8 assemblies. **(a)** XL-MS of AC8 shows self-association sites (homomultimeric links) (purple drops) evident of protein dimerisation. The links are on different sites of the protein that do not fit to only one dimeric structure but could be explained by multiple different dimeric structures. Regions covered in our AC8 structure are highlighted in grey. AC8 is copurified with CaM and Gαs (GNAS). Links with > 30 Å distance are considered violated crosslinks (red). Links with <30 Å distance are highlighted in blue. Intermolecular self-link sites are indicated as purple drops. The crosslinking map was produced with XiNET *(47)*. **(b-c)** The intermolecular crosslink positions in the AC8 structure are shown as purple spheres and lines, indicated on sketches. Oligomerisation sites are found on different AC8 domains, indicating that different AC8 complexes can exist. The juxtaposed models in c illustrate the crosslink positions. **(d-e)** Violated crosslinks are shown in the sketch (d) and indicated in the AC8 model (e). The violated crosslinks indicate that there is a lot of structural heterogeneity and flexibility in the sample. Violated crosslinks can also be a result of AC8 oligomerisation. For simplicity of the sketch, only selected violated crosslinks are highlighted, based on their domain location and consistent with the observed trend.

### XL-MS identifies the direct interaction sites of CaM, Gα_s_ and Gβγ

Our data on purified AC8 (**Fig. 5a**) show that the unresolved, flexible domains (N-terminus, C1b, C2b) of AC8 can be detected in close proximity to the structured domains. While many of the crosslinks can be explained with the cryo-EM structure of AC8 (**Fig. 5a**, blue crosslinks), some detected crosslinks between the C1b domain or the helical domain and the catalytic domains cannot be explained with our structure and could indicate an altered position of C1b or high domain flexibility (**Fig. 5c**, red crosslinks). The violated crosslinks (>30Å, violation cut-off based on the maximum Cα-Cα distance for DSS (*42*)) are highly reproducible and occur in all screened conditions (**Fig. S12**). We observed that Gαs and CaM were copurified with AC8 and were present in all conditions (**Fig. 5a, 6**).

The addition of CaM results in detection of crosslinks that were not detected when crosslinking AC8 alone. We find changes in the C1a and C2a domains but the links between C1a and C2a are not changed (**Fig. S13a**), suggesting that activation by CaM may cause conformational changes within the domains, with little or no change to the catalytic domain assembly.

XL-MS of AC8 in the presence of added Gαs (**Fig. 6b, f**) shows that the Gαs N-terminus is flexible and has multiple contact sites with the AC8 N-terminus, the C1a domain and the helical domain. The differences between the AC8-Gαs and the AC8-CaM-Gαs complex are only minute: analysis of the crosslinking data for these two samples shows the flexibility of the Gαs N-terminus. In the AC8-CaM-Gαs sample, we observe a crosslink between the Gαs N-terminus and CaM, suggesting that both can be in close proximity to each other. Additionally, we detect the same violated crosslink between AC8 and Gαs (Gαs E344xAC8 K581) in the AC8, the AC8-CaM, the AC8-CaM-Gαs and the AC8-Gβγ sample structure (**Figs. 6a-c, S13**). This indicates that Gαs may have several types of interactions in addition to the one resolved in the cryo-EM structure.

**Figure 6.**
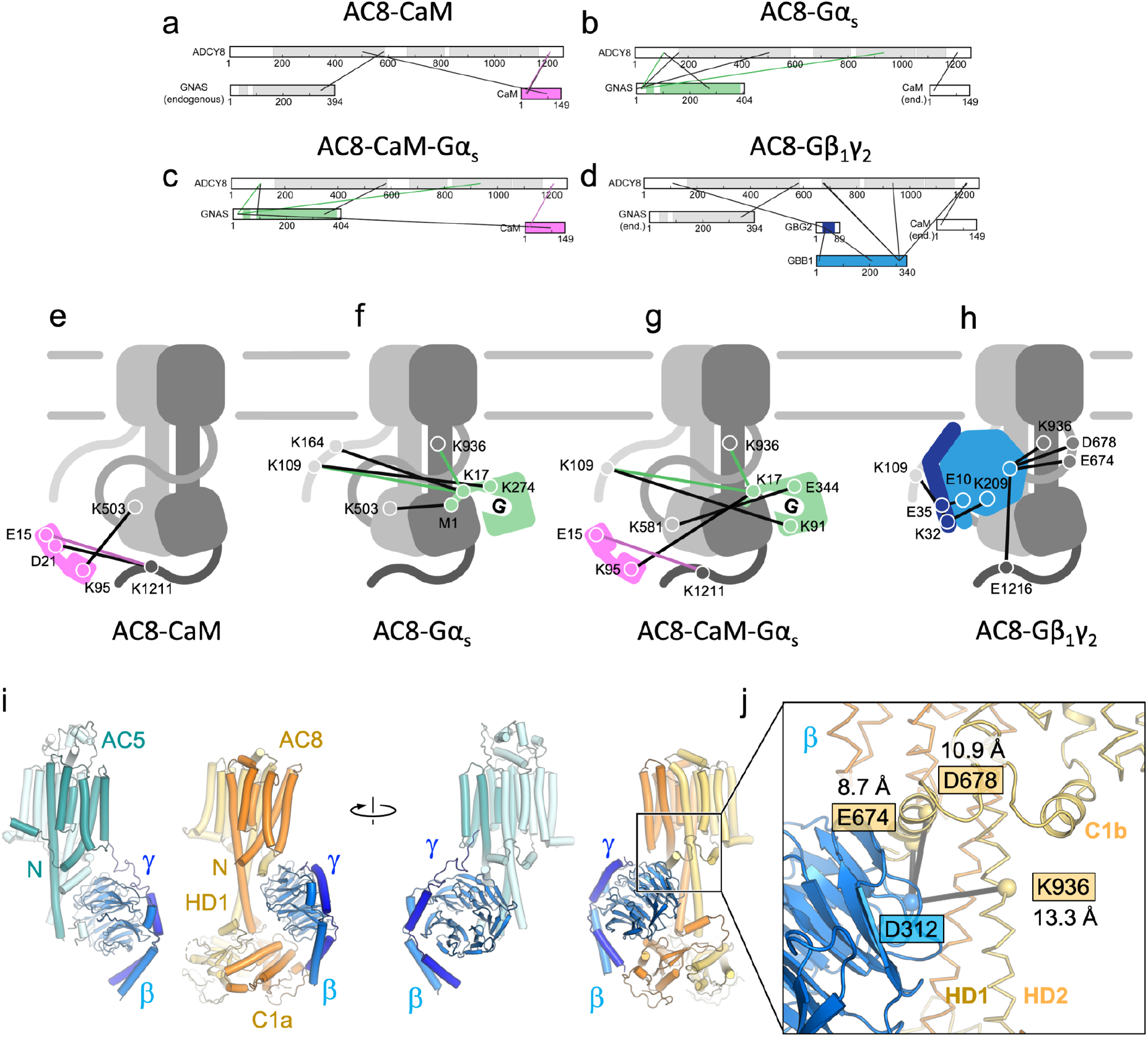
Crosslinks between AC8 and its interaction partners CaM, Gαs and Gβγ. **(a)** CaM interacts with the C2b domain of AC8 and is in close proximity to the C1a domain. Added interactors are coloured (CaM = pink, Gαs = green, Gβγ = blue), copurified interactors are labelled as “endogenous” or “end.”. Heteromeric crosslinks that were identified in at least one other condition are coloured (CaM = pink, Gαs = green, Gβγ = blue). **(b)** XL-MS analysis of AC8 with Gαs shows that the Gαs N-terminus has multiple contact sites with AC8. **(c)** XL-MS of AC8 with Gαs and CaM shows that CaM and Gαs are in close proximity with each other in a complex. **(d)** XL-MS of AC8 in the presence of Gβγ reveals a set of crosslinks sites between the Gγ and the N-terminus of AC8, and Gβ and the C1b and HD2 domains of AC8. (**e**,**f**,**g**) Sketches of AC8 and the added interactors, with the identified heteromeric crosslinks. Heteromeric crosslinks that were identified in at least one other condition are coloured (CaM = pink, Gαs = green, Gβγ = blue). **(h)** The crosslinking data facilitate protein docking, which reveals a plausible mode of Gβγ interaction with the AC8 Nt, C1b and C2b. The crosslinking maps were produced with XiNET *(47)*, docking was performed with HADDOCK *(48)*. Values in Å in white boxes indicate distances within the model of the docked AC8-Gβγ complex, consistent with the experimentally observed crosslinks. (i) Comparison between the AC5-Gβγ cryo-EM-based model (PDB ID: 8SL3) and our XL-MS and docking-based model of AC8-Gβγ complex reveals a remarkable similarity between the Gβγ binding sites in AC5 and AC8. (j) Residues of AC8 (yellow boxes) participating in crosslinking events with D312 of Gβγ (blue box) were used as docking restraints. The values in Å indicate the distances in the docking model.

In all analysed samples we could detect crosslinks of CaM with the C2b domain of AC8 (CaM E15xAC8 K1211). The AC8-CaM experiment also shows that CaM can be in close proximity to the AC8 C1a domain (CaM K95xAC8 K503) (**Fig. 6a, e**). The AC8-Gβγ experiment showed contact sites of Gβγ with the AC8 N-terminus (Gγ K32 x AC8 K109), the C1b domain (Gβ D312 x AC8 D674, Gβ D312 x AC8 D678), the helical domain (Gβ D312xAC8 K936) and the C2b domain (Gβ D312xAC8 E1216) (**Fig. 6h**). The crosslinks between the resolved parts of C1b, the helical domain and Gβγ facilitated protein docking and showed that Gβγ is positioned close to the helical domain, where it interacts with all three flexible domains (N-terminus, C1b, C2b) (**Fig. 6h**), further suggested by our LiP-MS experiment (**Fig. 4c**). When comparing the crosslinks detected in the AC8-Gβγ sample with the AC8 sample, we find additional crosslinks between the helical domain and the C1b domain (**Fig. S14d**). This finding supports the hypothesis that the interaction of AC8 with Gβγ causes a rearrangement between the helical domain and the C1b domain. Based on our data, we suggest that the interaction likely involves the AC8 N-terminus, C1b, the helical domain and the C2b domain. Interestingly, a recent cryo-EM study of the AC5-Gβγ complex (*43*) revealed an arrangement of the Gβγ subunits in the complex remarkably close to that observed in our docking model for AC8-Gβγ (**Fig. 6i-j**). As our attempts to perform cryo-EM analysis of AC8-Gβγ indicated that the complex may not be sufficiently ordered for direct structure determination, hybrid approaches are necessary to further characterize the modes of AC binding and regulation by Gβγ subunits.

## DISCUSSION

Our combined cryo-EM and structural proteomic approach applied to AC8 resulted in the following key findings (**Fig. 7**): (i) the cryo-EM structure of AC8 features a negatively-charged extracellular pocket, which may provide an interface for a yet unknown interaction partner; (ii) the forskolin-bound state of AC8 may not be conducive to tight nucleotide binding; (iii) AC8 is likely capable of forming oligomers, in a process that involves the flexible / unstructured protein domains, such as the N-terminus and the C1b domain; (iv) these flexible domains play a key role in the interactions between AC8 and its regulators (G proteins and CaM).

**Figure 7.**
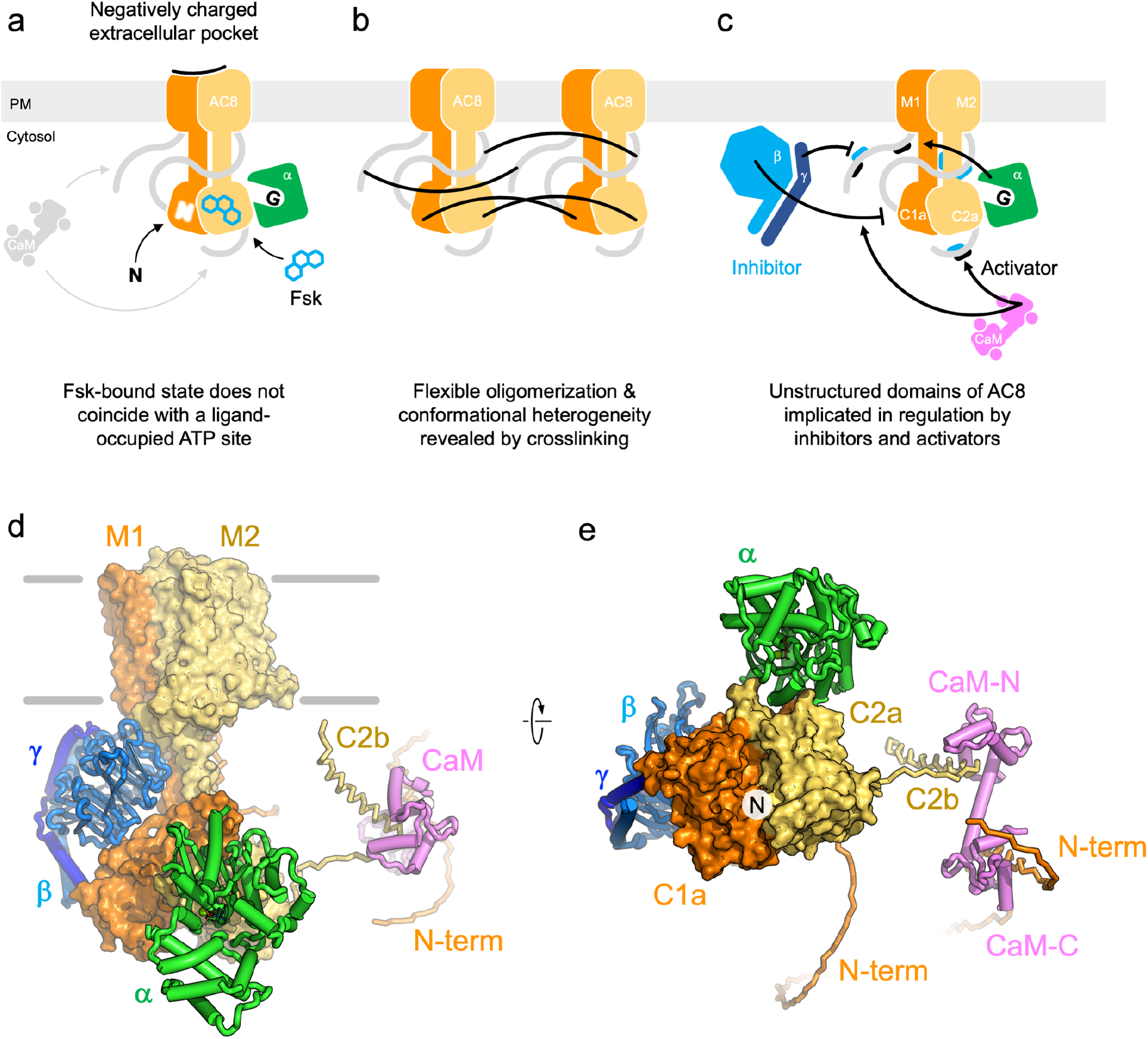
Insights into the structure of AC8 and interactions with its activators. **(a)** Cryo-EM structures of AC8 reveal the negatively charge extracellular pocket (red), and the dynamic nucleotide binding in presence of a small molecule activator, forskolin (Fsk). **(b)** XL-MS of purified AC8 provides direct evidence for its oligomerisation. AC8 oligomerisation involves the N-terminus and C1b domain **(c)** XL-MS and LiP-MS experiments show that the flexible N-, C-, and C1b domains of AC8 play vital roles in interactions with and regulation of AC8 by the activators (Gαs and CaM) and inhibitors (Gβγ). **(d-e)** An illustrative model of AC8 and its activating (Gαs and CaM) or inhibitory (Gβγ) interactors, based on cryo-EM (AC8-Gαs-CaM), docking (AC8-Gβγ) and AlphaFold2 (AC8-Gαs-CaM).

Contrary to previous studies on AC9 where obtaining a forskolin-bound state of the protein presented a substantial challenge (*31*), we observe a well-resolved density of forskolin bound to AC8, while the densities corresponding to the nucleotides (MANT-GTP or ATPαS), are less well defined. This is surprising, considering the ability of forskolin alone to fully stimulate AC8 cAMP production. The logical assumption we made prior to performing the experiments described here was that a potent activator, such as forskolin, should increase the affinity for the nucleotide. This should coincide with an improvement in the quality of the density corresponding to the bound nucleotide analogue. In contrast to this assumption, the 3D class that resulted in the best 3D reconstruction of AC8 features a well-resolved density for the bound forskolin and a poorly defined density corresponding to the active site (**Fig. 7a**). One possible interpretation of this finding is that forskolin-binding favours a conformation of AC8 that is compatible with high turnover rate (i.e., binding of ATP, fast conversion of ATP to cAMP, followed by rapid release of the products), but not with high affinity binding of the nucleotide analogues to the active site. It is conceivable that neither MANT-GTP nor ATPαS accurately recapitulate the interaction of ATP with the active site of AC8. We have previously observed the substantial flexibility of the AC9 ATP-binding site that can accommodate MANT-GTP in different poses dependent on the presence of forskolin (*32*). It is thus possible that AC8 binding of forskolin produces a conformation that does not favour a well-defined and uniform nucleotide pose. Furthermore, if a nucleotide-bound conformation of AC8 is not stable or is heterogeneous, the MANT-GTP- or ATPαS-bound states may be “lost” among the particles that are excluded during the 2D or 3D classification steps, as they do not contribute to a well-ordered protein conformation.

Cryo-EM analysis has become one of the primary tools for structure-function studies of proteins and protein complexes. The major caveat of this, and essentially every other structural biology approach, is that it relies on the conformational homogeneity of the sample. An ordered part of the protein can in many (if not most) cases be resolved by single particle analysis. This is the case for the relatively well-ordered portions of AC8: the TM region, the helical domain, the catalytic domains and the bound G protein subunit. Contrary to this, the unstructured regions of the AC8 present a major challenge for structural studies, as they cannot be resolved by conventional structural biology approaches, evident from the density map of AC8 that lacks the features corresponding to the N-terminus, the C1b and C2b domains. This challenge is not intrinsic to ACs alone since a wide range of proteins are known to be regulated by unstructured domains. However, specifically in the AC field interesting parallels can be drawn in the regulatory mechanisms of AC8 and the bacterial adenylyl cyclase toxins, *Bordetella pertussis* CyaA (*44*) and the *Bacillus anthracis* oedema factor (*45*). CaM has been found to interact with the disordered regions of these toxin ACs, allosterically communicating with the catalytic site and influencing the ordering of the catalytic site to stimulate the enzymatic activity. Careful computational and experimental analysis of the influence of CaM on the AC8 catalytic site dynamics will be necessary to establish whether similar modes of regulation apply to the mammalian Ca^2+^/CaM-sensitive ACs.

The combination of two structural proteomics approaches (LiP-MS and XL-MS) allowed us to overcome this inherent limitation and to (i) probe the conformational changes associated with regulator binding, and (ii) assess the candidate interaction sites and distances between AC8 and its regulators, deriving useful structural information. In addition to detecting the ability of AC8 to form oligomeric assemblies with a great degree of flexibility and/or heterogeneity (**Fig. 7b**), our data faithfully reproduce the previously observed CaM interactions at sites within the AC8 N-terminus and the C2b domain (**Fig. 7c**). To further illustrate the different modes of AC8 regulation by distinct binders, we complement our cryo-EM structure with the XL-MS-based docking model of AC8-Gβγ and the AlphaFold2 model of AC8-CaM-Gαs (**Fig. S16**). The AlphaFold2 model recapitulates the CaM binding sites in the N-terminus of AC8 and in the C2b domain, consistent with our data and with the published results (*25, 26*) remarkably well. This model helps us to rationalize the absence of a defined EM density for CaM in our cryo-EM maps, as the CaM-binding sites in AC8 are likely partially unstructured and detached from the ordered part of AC8 upon activation.

Our interpretations rely on the experimental evidence, cryo-EM analysis and AlphaFold2-based predictions. We believe that our cryo-EM-based reconstruction of the AC8-CaM-Gαs complex represents a CaM- and Gαs-bound complex wherein CaM and its binding sites can not be resolved by EM imaging. Nevertheless, it is worth noting that additional effects may affect the appearance of the particles under cryo-EM sample preparation and imaging conditions. For example, a biomolecular complex may prove to be unstable upon application to the cryo-EM grid, or upon exposure to the air-water interface. While we have no evidence that this is the case for the AC8-CaM-Gαs complex, due caution needs to be exercised particularly when inferring the presence of the components which can not be directly observed by imaging (i.e., CaM and the unstructured domains of AC8), even despite the available abundant evidence in support of their presence.

The combined illustrative model shown in **Fig. 7d**, along with our experimental results, provides the framework for understanding how Gαs, Gβγ and CaM engage AC8 via multiple interactions at the non-overlapping structured and unstructured regions, leading to activation or inhibition of cAMP production. While structural biology methods such as cryo-EM are not suitable for analysing the interactions of AC modulators with the unstructured regulatory sites of AC8 or other ACs, the MS-based approaches have shown their tremendous utility. The combination of cryo-EM, LiP-MS and XL-MS implemented here provides a powerful tool to study challenging membrane protein complexes that feature structured and unstructured domains.

## ACKNOWLEDGEMENTS

We thank Emiliya Poghosyan (EM Facility, PSI) and Miroslav Peterek (ScopeM, ETH Zurich) for expert support in cryo-EM data collection. We also thank Spencer Bliven and Marc Caubet-Serrabou (PSI) for the support in high performance computing. We thank Natalie de Souza and Jacopo Marino for valuable feedback on the manuscript. We thank the Wollscheid lab at ETH Zurich for allowing us to acquire data on their mass spectrometer. The work was supported by the Swiss National Science Foundation grant 184951 (VMK), and by a grant from the Vontobel foundation (VMK). P.P. is supported by the EPIC-XS Consortium (grant agreement no. 823839).

## Competing interests

Authors declare that they have no conflict of interest.

## Data and materials availability

The coordinates have been deposited in the Protein Data Bank, with entry codes 8BUZ, 8BV5. The cryo-EM density maps have been deposited in the Electron Microscopy Data Bank, with accession numbers EMD-16249, EMD-16252, EMD-16253, EMD-16254 and EMD-16255. The mass spectrometry proteomics data have been deposited to the ProteomeXchange Consortium via the PRIDE partner repository with the dataset identifiers PXD040303 (LiP-MS), PXD040374 (XL-MS) and PXD044766 (PRM). The R code used for data analysis is on GitHub (https://github.com/dschust-r/AC8_LiP_MS). All other data are available in the main text or the supplementary materials.

## Author contributions

B.K. planned and performed the experiments, analysed the data, wrote the manuscript, D.S. planned and performed the experiments, analysed the data, wrote the manuscript, M.O. and V.M. performed experiments, A.L. supervised XL-MS experiments, co-wrote the manuscript, P.P. supervised LiP-MS experiments, co-wrote the manuscript, V.M.K. planned and performed the experiments, analysed the data, wrote the manuscript.

## Notes

### Competing Interest Statement

The authors have declared no competing interest.

### Summary of Updates

This version of the manuscript features two sets of new experiments: (i) the FSEC-based AC8 / CaM-YFP binding assay; (ii) PRM- / MS-based quantitation of relative amounts of co-eluting CaM in the purified AC8 preparation. The manuscript includes a comparison between the AC8-Gbetagamma docking model (XL-MS-based) to the recently reported cryo-EM structure of AC5-Gbetagamma. Additionally, the manuscript includes an AlphaFold-based prediction of the AC8-Galphas-CaM model, consistent with the XL-MS and LiP-MS data. The manuscript features substantially revised illustrations, including an illustrative model of all AC8 modulators (G protein subunits and CaM), based on cryo-EM, docking and AlphaFold models.

## REFERENCES

1. K. F. Ostrom et al., Physiological roles of mammalian transmembrane adenylyl cyclase isoforms. Physiological Reviews 102, 815–857 (2022).

2. B. Khannpnavar, V. Mehta, C. Qi, V. Korkhov, Structure and function of adenylyl cyclases, key enzymes in cellular signaling. Current Opinion in Structural Biology 63, 34–41 (2020).

3. R. K. Sunahara, C. W. Dessauer, A. G. Gilman, Complexity and Diversity of Mammalian Adenylyl Cyclases. Annual Review of Pharmacology and Toxicology 36, 461–480 (1996).

4. N. Defer, M. Best-Belpomme, J. Hanoune, Tissue specificity and physiological relevance of various isoforms of adenylyl cyclase. American Journal of Physiology-Renal Physiology 279, F400–F416 (2000).

5. R. Iyengar, Molecular and functional diversity of mammalian Gs-stimulated adenylyl cyclases. The FASEB Journal 7, 768–775 (1993).

6. R. Taussig, J. A. Iñiguez-Lluhi, A. G. Gilman, Inhibition of Adenylyl Cyclase by Giα. Science 261, 218–221 (1993).

7. W.-J. Tang, A. G. Gilman, Type-Specific Regulation of Adenylyl Cyclase by G Protein βγ Subunits. Science 254, 1500–1503 (1991).

8. B. N. Gao, A. G. Gilman, Cloning and expression of a widely distributed (type IV) adenylyl cyclase. Proceedings of the National Academy of Sciences 88, 10178–10182 (1991).

9. M. L. Bayewitch et al., Inhibition of adenylyl cyclase isoforms V and VI by various Gβγ subunits. The FASEB Journal 12, 1019–1025 (1998).

10. X. Gao, R. Sadana, C. W. Dessauer, T. B. Patel, Conditional Stimulation of Type V and VI Adenylyl Cyclases by G Protein βγ Subunits*. Journal of Biological Chemistry 282, 294–302 (2007).

11. W. J. Tang, J. Krupinski, A. G. Gilman, Expression and characterization of calmodulin-activated (type I) adenylylcyclase. Journal of Biological Chemistry 266, 8595–8603 (1991).

12. D. Steiner, D. Saya, E. Schallmach, W. F. Simonds, Z. Vogel, Adenylyl cyclase type-VIII activity is regulated by Gβγ subunits. Cellular Signalling 18, 62–68 (2006).

13. S. Diel, K. Klass, B. Wittig, C. Kleuss, Gβγ Activation Site in Adenylyl Cyclase Type II: ADENYLYL CYCLASE TYPE III IS INHIBITED BY Gβγ*. Journal of Biological Chemistry 281, 288–294 (2006).

14. J. J. G. Tesmer et al., Two-Metal-Ion Catalysis in Adenylyl Cyclase. Science 285, 756–760 (1999).

15. J. J. Tesmer, R. K. Sunahara, A. G. Gilman, S. R. Sprang, Crystal structure of the catalytic domains of adenylyl cyclase in a complex with Gsalpha.GTPgammaS. Science 278, 1907–1916 (1997).

16. J. J. Tesmer, S. R. Sprang, The structure, catalytic mechanism and regulation of adenylyl cyclase. Curr Opin Struct Biol 8, 713–719 (1998).

17. J. Chen et al., A Region of Adenylyl Cyclase 2 Critical for Regulation by G Protein βγ Subunits. Science 268, 1166–1169 (1995).

18. C. Wittpoth, K. Scholich, Y. Yigzaw, T. M. Stringfield, T. B. Patel, Regions on adenylyl cyclase that are necessary for inhibition of activity by βγ and Giα subunits of heterotrimeric G proteins. Proceedings of the National Academy of Sciences 96, 9551–9556 (1999).

19. C. S. Brand, R. Sadana, S. Malik, A. V. Smrcka, C. W. Dessauer, Adenylyl Cyclase 5 Regulation by Gβγ Involves Isoform-Specific Use of Multiple Interaction Sites. Molecular Pharmacology 88, 758 (2015).

20. K. R. Westcott, D. C. La Porte, D. R. Storm, Resolution of adenylate cyclase sensitive and insensitive to Ca2+ and calcium-dependent regulatory protein (CDR) by CDR-sepharose affinity chromatography. Proceedings of the National Academy of Sciences 76, 204–208 (1979).

21. E.-J. Choi, Z. Xia, D. R. Storm, Stimulation of the type III olfactory adenylyl cyclase by calcium and calmodulin. Biochemistry 31, 6492–6498 (1992).

22. J. Sanchez-Collado, J. J. Lopez, I. Jardin, G. M. Salido, J. A. Rosado, in Reviews of Physiology, Biochemistry and Pharmacology, S. H. F. Pedersen, Ed. (Springer International Publishing, Cham, 2021), pp. 73–116.

23. M. J. Berridge, P. Lipp, M. D. Bootman, The versatility and universality of calcium signalling. Nature Reviews Molecular Cell Biology 1, 11–21 (2000).

24. A. Raffaello, C. Mammucari, G. Gherardi, R. Rizzuto, Calcium at the Center of Cell Signaling: Interplay between Endoplasmic Reticulum, Mitochondria, and Lysosomes. Trends in Biochemical Sciences 41, 1035–1049 (2016).

25. N. Masada, A. Ciruela, D. A. MacDougall, D. M. F. Cooper, Distinct Mechanisms of Regulation by Ca2+/Calmodulin of Type 1 and 8 Adenylyl Cyclases Support Their Different Physiological Roles*. Journal of Biological Chemistry 284, 4451–4463 (2009).

26. S. Herbst et al., Structural insights into calmodulin/adenylyl cyclase 8 interaction. Analytical and Bioanalytical Chemistry 405, 9333–9342 (2013).

27. R. E. Simpson, A. Ciruela, D. M. Cooper, The role of calmodulin recruitment in Ca2+ stimulation of adenylyl cyclase type 8. J Biol Chem 281, 17379–17389 (2006).

28. K. E. Smith, C. Gu, K. A. Fagan, B. Hu, D. M. Cooper, Residence of adenylyl cyclase type 8 in caveolae is necessary but not sufficient for regulation by capacitative Ca(2+) entry. J Biol Chem 277, 6025–6031 (2002).

29. C. Gu, D. M. F. Cooper, Calmodulin-binding Sites on Adenylyl Cyclase Type VIII. Journal of Biological Chemistry 274, 8012–8021 (1999).

30. D. A. MacDougall, S. Wachten, A. Ciruela, A. Sinz, D. M. F. Cooper, Separate Elements within a Single IQ-like Motif in Adenylyl Cyclase Type 8 Impart Ca2+ Calmodulin Binding and Autoinhibition Journal of Biological Chemistry 284, 15573–15588 (2009).

31. C. Qi, S. Sorrentino, O. Medalia, V. M. Korkhov, The structure of a membrane adenylyl cyclase bound to an activated stimulatory G protein. Science 364, 389–394 (2019).

32. C. Qi et al., Structural basis of adenylyl cyclase 9 activation. Nature Communications 13, 1045 (2022).

33. M. Erdorf, T.-C. Mou, R. Seifert, Impact of divalent metal ions on regulation of adenylyl cyclase isoforms by forskolin analogs. Biochemical Pharmacology 82, 1673–1681 (2011).

34. A. Seth et al., Distinct glycerophospholipids potentiate Gsα-activated adenylyl cyclase activity. Cellular Signalling 97, 110396 (2022).

35. T. A. Baldwin, Y. Li, C. S. Brand, V. J. Watts, C. W. Dessauer, Insights into the Regulatory Properties of Human Adenylyl Cyclase Type 9. Molecular Pharmacology 95, 349 (2019).

36. V. Mehta et al., Structure of Mycobacterium tuberculosis Cya, an evolutionary ancestor of the mammalian membrane adenylyl cyclases. eLife 11, e77032 (2022).

37. A. Holfeld et al., Systematic identification of structure-specific protein–protein interactions. bioRxiv, 2023.2002.2001.522707 (2023).

38. A. Leitner, T. Walzthoeni, R. Aebersold, Lysine-specific chemical cross-linking of protein complexes and identification of cross-linking sites using LC-MS/MS and the xQuest/xProphet software pipeline. Nat Protoc 9, 120–137 (2014).

39. A. Leitner et al., Chemical cross-linking/mass spectrometry targeting acidic residues in proteins and protein complexes. Proc Natl Acad Sci U S A 111, 9455–9460 (2014).

40. C. Gu, J. J. Cali, D. M. Cooper, Dimerization of mammalian adenylate cyclases. Eur J Biochem 269, 413–421 (2002).

41. W. J. Tang, M. Stanzel, A. G. Gilman, Truncation and alanine-scanning mutants of type I adenylyl cyclase. Biochemistry 34, 14563–14572 (1995).

42. E. D. Merkley et al., Distance restraints from crosslinking mass spectrometry: mining a molecular dynamics simulation database to evaluate lysine-lysine distances. Protein Sci 23, 747–759 (2014).

43. C. L. Chen et al., Molecular basis for Gbetagamma-mediated activation of phosphoinositide 3-kinase gamma. bioRxiv, (2023).

44. D. P. O’Brien et al., Calmodulin fishing with a structurally disordered bait triggers CyaA catalysis. PLoS Biol 15, e2004486 (2017).

45. C. L. Drum et al., Structural basis for the activation of anthrax adenylyl cyclase exotoxin by calmodulin. Nature 415, 396–402 (2002).

46. H. Ashkenazy et al., ConSurf 2016: an improved methodology to estimate and visualize evolutionary conservation in macromolecules. Nucleic Acids Research 44, W344–W350 (2016).

47. C. W. Combe, L. Fischer, J. Rappsilber, xiNET: cross-link network maps with residue resolution. Mol Cell Proteomics 14, 1137–1147 (2015).

48. G. C. van Zundert, A. M. Bonvin, Modeling protein-protein complexes using the HADDOCK webserver “modeling protein complexes with HADDOCK”. Methods Mol Biol 1137, 163–179 (2014).

