## Supplementary information for "Regulatory sites of CaM-sensitive adenylyl cyclase AC8 revealed by cryo-EM and structural proteomics"

**Affiliations:**

### MATERIALS AND METHODS

#### *Chemicals*

Detergents, dodecyl-β-maltoside (DDM), glyco-diosgenin (GDN), digitonin, cholesteryl hemisuccinate (CHS) and Brain Polar Lipids were purchased from Antrace Inc. Calmodulin (from bovine testes, Catalog No: P1431) was purchased from Sigma-Aldrich. All other chemicals were obtained from Sigma-Aldrich (St. Louis, MO, USA), unless indicated otherwise.

#### *Protein expression and purification*

*Adenylyl cyclase AC8:* The full-length bovine AC8 (Uniprot: E1BQ12) was cloned into pACMV-based tetracycline-inducible vector with C-terminal 3C-YFP-TwinStrep fusion tag. The plasmids were transfected into HEK 293F GnTI- cells using branched polyethyleneimine (PEI) and a stable monoclonal cell line (HEK293-AC8) capable of expressing AC8 was generated for large scale expression. HEK293-AC8 cells were cultured in Protein Expression Medium (PEM, Gibco) to a cell density of $1.8\times10^{6}$ $ml^{-1}$ and expression was induced by adding 200 μL of 10 mg/mL Tetracycline. For purification, the cell pellets were resuspended in 50 mM Tris-HCl pH 8.0, 150 mM NaCl, 10 % glycerol supplemented with protease inhibitors (1 mM benzamidine, 1 μg/ml leupeptin, 1 μg/ml aprotinin, 1 μg/ml pepstatin, 1 μg/ml trypsin inhibitor and 1 mM PMSF). Cells were lysed using a Dounce homogenizer and the total cell membranes were collected by ultracentrifugation (Ti45 rotor, 186,000 x g for 40 min at 4 °C). Membranes were flash-frozen and stored at -80°C until needed the day of experiment. On the day of purification, membranes were thawed, and resuspended in the 50 mM Tris-HCl pH 8.0, 150 mM NaCl, 10 % Glycerol, 1 % DDM and 0.02 % cholesteryl hemisuccinate (CHS) and solubilized for 1 h at 4°C. The detergent solubilized lysate was cleared by ultracentrifugation (Ti45 rotor, 186,000 x g for 40 min at 4 °C) to remove insoluble debris. The supernatant was incubated with 4 mL of CNBr-activated Sepharose coupled to an anti-GFP nanobody. After a 60 min incubation at 4 °C the resin was collected in a gravity column and washed with 40 column volumes of 50 mM Tris-HCl pH 8.0, 150 mM NaCl, 0.02 % GDN, 5 % glycerol. The protein was eluted using cleavage by HRV 3C protease (1:10 w/w). The eluted protein was concentrated with a 100 kDa cut-off Amicon Ultraspin device (Millipore) and subjected to size-exclusion chromatography (SEC) using a Superpose 6 Increase 10/300 GL column (GE Healthcare) equilibrated in 50 mM Tris-HCl pH 8.0, 150 mM NaCl, 0.02 % GDN. The fractions corresponding to purified AC8 (elution volume, 13-16 mL) were collected. Finally, SEC fractions were concentrated and used directly for cryo-EM grids preparation or flash-frozen in 10 % glycerol and stored at -80 °C. The protein purity was assessed using 4–20 % SDS-PAGE (BIO-RAD) and visualized by standard Coomassie brilliant blue staining technique. Protein concentration was determined by absorbance at 280 nm using $\varepsilon = 126975\ M^{-1}\ cm^{-1}$.

*G protein $\alpha s$ subunit***.** The procedure for expression and purification of the bovine Gαs (Uniprot ID P04896-1, with a C-terminal 8xHis-tag) was similar to the one previously described (*1, 2*). The standard Bac-to-Bac baculovirus expression system was used (Invitrogen): High Five (Hi5) insect cells were cultured in suspension to a cell density of $1.5\times10^{6}$ $ml^{-1}$ and 1-2 % P2 virus was added to infect the cells for protein expression.

The cells were harvested after 72 hours. For protein purification, the cells were resuspended in buffer A, lysed using a Dounce homogenizer and solubilized by adding 1% DDM for 1 hour. The lysate was clarified by ultracentrifugation (Ti45 rotor, 186000 x g for 40 min at 4 °C). The supernatant was incubated with 1 ml Ni-NTA resin for 30 min. The resin was washed with 50 mM Tris-HCl pH 8.0, 150 mM NaCl, 0.02 % DDM, and 20 mM imidazole. A second wash followed this with 50 mM Tris-HCl pH 8.0, 150 mM NaCl, 0.02 % GDN, 5 % glycerol, 40 mM imidazole. The protein was eluted with 50 mM Tris-HCl pH 8.0, 150 mM NaCl, 0.02 % GDN, 5 % glycerol, 250 mM imidazole. The eluted protein was concentrated with a 30-kDa cut-off Amicon Ultraspin device (Millipore) and injected on Superose 6 Increase 10/300 GL column (GE Healthcare) pre-equilibrated in 50 mM Tris-HCl pH 8.0, 150 mM NaCl, 0.02 % GDN. The peak fractions (elution volume, 14-16 mL) were concentrated, snap-frozen in liquid nitrogen in aliquots, and stored at -80 °C until the day of the experiment. Protein concentration was determined by absorbance at 280 nm using $\varepsilon = 43360\ M^{-1}\ cm^{-1}$.

*G protein beta gamma subunit:* Gβγ was expressed from insect cells (Hi5 cells) using flashBAC system(*3*). In general, Hi5 insect cells were cultured in serum free SF-4 medium to a cell density of $1.5\times10^{6}\ ml^{-1}$ and then expression of protein was induced by adding 1-2 % P2 virus. After 48 h cells were harvested by centrifugation at 1,000 x g for 15 minutes. Cell pellets were resuspended in buffer A, lysed using a Dounce homogenizer and the total cell membranes were collected by ultracentrifugation (Ti45 rotor, 186000 x g for 40 min at 4 °C). The supernatant was discarded and the membrane pellet was resuspended in 50 mM Tris-HCl (pH 8), 200 mM NaCl, 1 % sodium cholate, 10 % glycerol, 25 mM imidazole and left rotating for 1 hour at 4°C. The suspension was clarified using ultracentrifugation centrifuged at 40,000 rpm (186,000 x g; Beckman Ti45 rotor) for 40 minutes at 4 °C and the supernatant was incubated with 2 mL HisPure cobalt resin (ThermoFisher Scientific) per 1 L original culture for 1 hour at 4 °C under constant agitation. Bead/sample suspension was loaded onto a gravity column and washed with 40 column volumes of 50 mM Tris pH 8.0, 150 mM NaCl, 10 % glycerol, 25 mM imidazole and 0.1 % digitonin, followed by washing with 40 column volumes of 50 mM Tris pH 8.0, 150 mM NaCl, 10 % glycerol, 50 mM imidazole and 0.1 % digitonin). Gβγ was eluted with 50 mM Tris pH 8.0, 150 mM NaCl, 10 % glycerol, 400 mM imidazole and 0.1 % digitonin and the His-tag was cleaved over night with HRV 3C protease (amount: 1:50 eluted protein:HRV 3C). The flow-through was collected and desalted with buffer (50 mM Tris pH 8.0, 150 mM NaCl, 10 % glycerol and 0.1 % digitonin) using a GE PD-10 desalting column. The desalted protein was then passed through Ni-NTA resin to remove HRV 3C protease, concentrated to 1 mL and injected on Superose 6 Increase 10/300 GL column (GE Healthcare) equilibrated in 50 mM Tris-HCl, 150 mM NaCl, 0.02 % GDN, pH 8.0. Fractions (elution volume, 13-16 mL) containing Gβγ were further concentrated and flash frozen in liquid nitrogen with the addition of 10 % glycerol and stored at -80 °C until further use.

*GFP-nanobody*: The expression and purification of anti GFP nanobody was carried out as previously described (*4*). In short, the anti-GFP nanobody plasmid was transformed into E. coli BL21 (DE3) and grown in Luria Broth (LB) media supplemented with 50 µg/mL ampicillin at 37°C. Once the optical density at 600 nm (OD600) of bacterial culture reached 0.5, the expression of protein was induced by adding 0.5 mM IPTG followed by overnight incubation at the 20 °C. Bacterial cultures were harvested at 4,000 g for 20 min at 4 °C, and pellets were frozen in liquid N2 and stored at −80 °C. Frozen pellets were thawed on ice, re-suspended in 25 mM HEPES pH 8.0, 150 mM NaCl, 10 mM Imidazole, 1 mM PMSF, 10 µg/mL DNase I. The cells were lysed by sonication, and the cell lysate was centrifuged for 30 min at 20,000 x g. Clarified lysate was then incubated with Ni-NTA resin (1-2 mL bed volume of resin per 1 L of culture) for 30-40 minutes, and subsequently washed with 20 CV of 25 mM HEPES pH 8.0, 150 mM NaCl, 50 mM imidazole, and later protein was eluted with 5 CV of same buffer with 250 mM Imidazole. The elution was concentrated with 10 kDa cut-off Amicon concentrator and loaded on a Superdex 75 16/600 GL column (GE Healthcare) in 25 mM HEPES pH 8.0, 150 mM NaCl. SEC fraction corresponding to GFP-nanobody was pooled and flash frozen in liquid nitrogen, and stored at -80 °C. Protein concentration was determined by absorbance at 280 nm using ε = 27055 $M^{-1}$ $cm^{-1}$.

*Membrane Scaffold Protein:* The expression and purification of Membrane Scaffold Protein MSP1E3D1 was carried out as previously described(*5*). In brief, the MSP1E3D1 plasmid was transformed into E. coli BL21 (DE3) and grown at 37 °C in Terrific Broth (TB) media. The expression of protein was induced with 1 mM IPTG at OD600 of ~2-3 and incubation for 3 h. After harvesting by centrifugation, cell pellets were re-suspended in lysis buffer (50 mM Tris-HCl pH 8.0, 200 mM NaCl, 25 mM imidazole, 1 % Triton-X100, 1mM PMSF and 10 µg/ml DNaseI) and cells were lysed by sonication. The clarified lysate after centrifugation (20,000 x g for 30 minutes) was incubated with Ni-NTA resin for 30 min, and subsequently step washed with 10 CV of buffer (50 mM Tris-HCl pH 8.0, 150 mM NaCl, 25 mM imidazole, 1 % Triton-X100), 5 CV of buffer (50 mM Tris-HCl pH 8.0, 150 mM NaCl, 25 mM imidazole, 2 % Sodium Cholate), and 5 CV of buffer (50 mM Tris-HCl pH 8.0, 150 mM NaCl, 50 mM Imidazole). Finally, the protein was eluted in 50 mM Tris-HCl pH 8.0, 150 mM NaCl, 350 mM imidazole. Elution fractions containing MSP1E3D1 were pooled, desalted in buffer (20 mM Tris-HCl pH 8.0, 200 mM NaCl) and flash frozen in liquid nitrogen, and stored at -80 °C. Protein concentration was determined by absorbance at 280 nm using ε = 29,400 $M^{-1}$ $cm^{-1}$.

*Calmodulin-YFP:* The HEK293F cells grown in suspension at a cell density of 1.5 million cells/ml of culture were transfected with an expression plasmid containing the CaM-YFP-twinStrep (referred to here and throughout the manuscript as CaM-YFP) construct (1 mg DNA per 1 L of cells, using linear PEI at a DNA to PEI ratio of 1:3). The cells were incubated in a shaker for 72 h at 37 °C, at 5 % $CO_2$ and 120 rpm, after which the cell pellets were harvested by centrifugation at 2000 rpm at 4 °C for 20 min using a 6 x 1000 mL swinging bucket rotor (Sorvall BIOS 16 Centrifuge, Thermo Scientific). The collected cell pellets were frozen and stored at -80 °C until the day of purification. To purify CaM-YFP, the cell pellets corresponding to 1 L of culture were thawed, resuspended in a cold buffer (50 mM Tris-HCl, pH 7.5, 200 mM NaCl, 2 mM EGTA, 1 µg/µL DNase I, 1x Roche Complete EDTA-free protease inhibitor cocktail) and lysed using a Dounce homogenizer. The lysate was clarified by ultracentrifugation (Ti45 rotor, 186,000 x g for 40 min at 4 °C), and the supernatant was incubated with 4 ml Strep-Tactin® Superflow® resin for 30 min at 4 °C with rotation. After incubation the sample was applied to a gravity column and washed with 40 column volumes of buffer (50 mM Tris-HCl, pH 7.5, 200 mM NaCl, 2 mM EGTA). The protein was eluted with 5 column volumes of buffer (50 mM Tris-HCl, pH 7.5, 200 mM NaCl, 2 mM EGTA) containing 5 mM desthiobiotin. The sample was concentrated with a 10 kDa cut-off AmiconUltra® concentrator to a final volume of 1 ml, and further purified by HPLC using a Superdex 200 10/300 GL column pre-equilibrated with a buffer (50 mM Tris-HCl, pH 7.5, 200 mM NaCl, 2 mM EGTA) containing additionally 10% glycerol. The fractions corresponding to CaM-YFP were pooled, concentrated, aliquoted, flash frozen in liquid nitrogen, and stored at -80 °C until the day of experiment.

***Parallel reaction monitoring (PRM) –based quantification of AC8 and CaM***

*Sample preparation:* 2 aliquots of purified AC8 (10 µg) at a concentration of 0.23 mg/mL were digested with protiFi S-Trap™ micro columns according to the manufacturer's protocol. After drying in a vacuum centrifuge the peptide mixture was resuspended in 50 µL 5% acetonitrile (ACN), 0.1% formic acid (FA) in water.

Heavy labeled AQUA peptides (4 peptides for AC8, 3 peptides for CaM; listed in Table S1) were custom synthesized and ordered from Thermo Fisher Scientific. A bovine serum albumin (BSA) background mix containing 100 fmol/µL BSA peptides, iRT peptides (Biognosys), 5% ACN and 0.1% FA in water was prepared. The AQUA peptides were mixed and diluted in 1:10 steps to obtain the calibration curve samples. Briefly, for the 500 fmol/µL sample 2 µL of each peptide stock (5 pmol/µL) were mixed and 6 µL of the BSA background mix were added (20 µL final volume). This sample was sequentially diluted in 1:10 dilution steps (2 µL sample + 18 µL BSA background) to obtain samples containing 50 fmol/µL, 5 fmol/µL, 500 amol/µL, 50 amol/µL, 5 amol/µL and 500 zmol/µL of each peptide.

Purified samples were prepared by diluting the samples 1:4 in 5% ACN, 0.1% FA, with the addition of AC8 AQUA peptides at ca. 500 fmol/µL and CaM AQUA peptides at ca. 500 amol/µL, iRT peptides and BSA background.

*LC-MS/MS – PRM data acquisition:* The samples were analyzed on an Exploris 480 mass spectrometer (ThermoFisher Scientific) connected to a Vanquish Neo (ThermoFisher Scientific) UHPLC. In separate runs, 0.5 µL and 1 µL of the samples were injected and peptides were separated on a 40 cm x 0.75 µm (inner diameter) column in-house packed with 1.9 µm C18 beads (Dr. Maisch Reprosil-Pur 12) over a 30 minute linear gradient from 7-35% B (A: 0.1% FA, B: 80% ACN, 0.1% FA) at 300 nL/min, at 50°C. MS1 spectra were acquired within a scan range between 150-2000 m/z at an Orbitrap resolution of 30,000. RF lens was set to 50%, the AGC target was set to "Standard". The selected precursors (including iRT peptides) and settings are listed in Table S1.

*Data analysis:* The acquired data was analyzed in Skyline (v. 23.0.9.187), exported and further interpreted in R using the R packages tidyverse and data.table. The retention times of the detected iRT peptides were checked for consistency. To produce the calibration curve, the calculated areas of the 5 highest ranked fragments of heavy peptides were summed up for each sample. After assessing linearity of the calibration curve (LOQ) (Fig. S2), the two lowest concentration points (500 zmol/µL and 5 amol/µL) were removed. A linear model (lm) was fit onto the log10 transformed, summed up area values at the specific log10 transformed concentration points (log10(area) ~ log10(concentration)) and the slope and intercept were calculated for each heavy labeled AQUA peptide.

To assess AC8 and CaM sample concentrations and their stoichiometry the 5 highest ranked fragments of light peptides were summed up for each sample. The slope and intercept values obtained from the respective calibration curves were used to calculate the absolute concentrations of peptides and proteins (average of peptide intensity). Peptide EAFSLFDK (CaM) was excluded from the calculations, since the measurement points were outside of the range of linearity. The average injected protein amount of AC8 for 0.5 µL injection volume was calculated to be 60.69 ± 11.39 fmol (mean ± standard error of the mean), for 1 µL 113.58 ± 21.03 fmol. The average amount of CaM for 0.5 µL injection volume was 0.17 ± 0.07 fmol, for 1 µL 0.33 ± 0.12 fmol. The ratio of CaM:AC8 in the purified protein samples was in the range of 1:200 – 1:720. CaM can therefore be estimated to be co-purified with AC8 in substoichiometric amounts at an approximate ratio of 1:500 (CaM:AC8).

#### *Reconstitution of AC8 into nanodiscs*

Reconstitution of AC8 in MSP1E3D1 nanodiscs was performed using freshly purified protein. In brief, 2.5 mg of brain polar lipid (BPL) dissolved in 100 µL of chloroform was dried under a stream of nitrogen. The dried film of lipid was mixed with 300 µL of 3 % DDM and the lipid-detergent mixture was then sonicated using bath sonicator (Bandelin SONOREX™SUPER, Germany) until the mixture turned translucent. Detergent-solubilized BPL extract was added to freshly purified AC8 at a molar ratio of 1:150, and incubated for 30 min at room temperature with gentle rotation. MSP1E3D1 was added to the protein-lipid mixture and incubated for additional 30 min. Concentrations of AC8 for nanodisc reconstitution were in the 2-3 µM range, with a molar ratio of AC8 to MSP1E3D1 to lipid of 1:2:150. Following the incubation period, nanodisc formation was triggered by adding 150 mg of wet Bio-beads (washed with 100 % methanol and with Milli-Q water). This final reconstitution mixture was incubated at 4 °C for 16 h (overnight) with gentle mixing. The supernatant was cleared of the beads by letting the beads settle and removing liquid carefully with a pipette. Sample was spun for 10 min at 25000 x g using a bench-top Eppendorf centrifuge before loading onto a Superose 6 Increase 10/300GL column equilibrated in 20 mM Tris-HCl pH 8.0, 150 mM NaCl. The peak fractions corresponding to AC8 in MSP1E3D1 (elution volume, 13-15.2 mL) were collected, concentrated with a 100 kDa cutoff Amicon concentrator and used for cryo-EM grid preparation.

#### *Adenylyl cyclase activity assays*

The cAMP accumulation assay was performed using protocol adapted from Alvarez and Daniels (*1, 2, 6*). Briefly, the purified AC8 was diluted to 20-30 nM in buffer A (50 mM Tris PH 7.5, NaCl, 0.02% GDN) for all the experiments. Similarly, AC8 complexes with varying concentration of CaM , Gαs, Gβγ and forskolin were prepared. The enzymatic reaction was then initiated by mixing 100 μL of protein solution with 100 μL of 2X reaction mixture containing 4 mM $MgCl_2$, 10 mM $MnCl_2$, 0.2 mM ATP, and 20 nM of $H^3$-ATP (PerkinElmer). The reaction mixture was then incubated at 30 °C for 30 min. After incubation, the reaction was then quenched by the addition of 20 μL of 2.2 M HCl for 4 min at 95 °C. The quenched reaction mixture was immediately cooled on ice, followed by loading onto gravity columns packed with 1.3 g of packed aluminium oxide. cAMP which binds less strongly than ATP, was eluted with 4 ml of 0.2 M ammonium acetate. Before counting, 12 mL of scintillation liquid (LabLogic) was added, and the final solution was brought to counting on a Packard 2250 CA Tri Carb liquid scintillation counter.

***CaM binding assays***

AC8-CaM interaction was analyzed using fluorescence size exclusion chromatography (FSEC). For this purpose the purified CaM-YFP was used as a fluorescent probe and binding of CaM to AC8 was detected based on the emergence of YFP fluorescence at the position of the eluted AC8 peak in an analytical SEC experiment. In brief, purified tag-free AC8 (100 nM final concentration) was mixed with various amounts of CaM-YFP (at a range of concentrations, from 3.75 to 1000 nM) in a final volume of 100 μl. The mixtures were incubated on ice for 15 min, in the presence of $CaCl_2$ (1 mM) or EGTA (1 mM). After the incubation, each sample was injected onto the Agilent SEC5-300 column (flow 0.3 ml/min, excitation 488 nm, emission 510 nm) pre-equilibrated in 50 mM Tris HCl, pH 8.0, 150 mM NaCl, 0.02% GDN and 1 mM $CaCl_2$ or 1 mM EGTA. The total peak area (elution volume, 1.2-1.62 ml) and peak intensity (at 1.47 ml) were quantified for each experiment. The binding affinity of AC8 and CaM was evaluated using the one-site saturation binding model in GraphPad Prism 8. Co-localisation of CaM-YFP with AC8 in the complex peak was validated by analysing the full AC8-CaM-YFP complex (1:2 ratio) by SEC, using Superose 6 increase column, followed by SDS-PAGE, in-gel fluorescence (using BIO-RAD ChemiDoc™ MP Imaging System device) and Coomassie blue staining.

***Cryo-EM sample preparation and data collection***

A typical sample of AC8-CaM-Gαs complex was prepared by mixing concentrated AC8 (~6 mg/mL) with 1.2-fold molar excess of GTPγS-activated Gαs protein, two fold excess of calmodulin (purchased from Sigma-Aldrich, Catalog No: P1431) and incubated with 0.5 mM MANT-GTP, 0.1mM GTPγS, 1 mM $CaCl_2$, 2 mM $MgCl_2$, and 5 mM $MnCl_2$. The final concentration of AC8 in the sample was ~5 mg/ml. The final Quantifoil 1.2/1.3 200-mesh grids were briefly glow discharged in a PELCO easiGlow (Ted Pella) glow discharge cleaning system for 25 s at 30 mA in air. A 3.5 μL sample was deposited on the carbon side of the grid and blotted for 3 s (blot force of 20) inside the chamber of a Vitrobot Mark IV (Thermo Fisher Scientific) with 100 % humidity and at 4 °C. The grid was then plunged into liquid ethane, and the grids were stored in liquid nitrogen. A total of three datasets with 3618, 5178, and 7149 movies were collected using EPU on a 300 kV Titan Krios (Thermo Fisher Scientific) equipped with a Gatan K3 direct electron detector and a Gatan Quantum-LS GIF at ScopeM, ETH Zurich. All movies were acquired in super-resolution mode with a defocus range of

−0.5 to −3 µm and a final calibrated pixel size of 0.33 Å. The total dose per movie was 60, 56 and 49 $e^-/Å^2$ for datasets 1, 2, and 3, respectively.

For AC8-Gαs-$Ca^{2+}$/CaM complex in lipid nanodiscs, the AC8 reconstituted in lipid nanodisc at a concentration of 4 mg/ml was incubated with 1.2-fold molar excess of GTPγS -activated Gαs protein, two fold excess of calmodulin 1 mM ATPαs, 0.1 mM GTPγS, 1 mM $CaCl_2$, 2 mM $MgCl_2$, and 5 mM $MnCl_2$ for 10 min at 4 °C. A small aliquot of the sample (3.5 µl) was applied to the glow-discharged Quantifoil R1.2/1.3 200-mesh grid. The grid was blotted for 3 s and plunge-frozen in the liquid ethane using a Vitrobot Mark IV (Thermo Fisher Scientific). The grids were transferred to and stored in liquid nitrogen for subsequent cryo-EM data collection. A total of three datasets with 2007, 8022, and 11291 movies were collected using EPU on a 300 kV Titan Krios (Thermo Fisher Scientific) equipped with a Gatan K3 direct electron detector and a Gatan Quantum-LS GIF at ScopeM, ETH Zurich. All movies were acquired in super-resolution mode with a defocus range of −0.5 to −3 µm and two-fold binned with final calibrated pixel size of 0.66 Å. Movies of 40 frames were collected with a total dose of 55, 55, and 56.4 $e^-/Å^2$ for datasets 1, 2, and 3, respectively.

#### *Cryo-EM data analysis and model building*

The data processing of AC8-CaM-Gαs was performed in Relion (3.0 and 3.1-beta versions) (*7*). All movie stacks were motion corrected using MotionCorr 1.1.0 (*8*) and binned twofold to yield a magnified pixel size of 0.66. All micrographs were CTF corrected using GCTF (*9*). Initially, around 1017 particles were manually picked to generate 2D template for autopicking. Selected 2D classes from manually picked particles were used to autopick particles from all micrographs in dataset 1. After multiple rounds of 2D and 3D classification jobs, the best 3D class from first dataset was used to repick the particle with 3D projections for all three of AC8-CaM-Gαs complex datasets. 5.72 million particles were picked from three datasets. After multiple rounds of 2D and 3D clarifications, a set of 615431 particles was selected for masked 3D refinement (using masks including or excluding the nanodisc density), resulting in a 3D reconstruction at 4.5 Å. The refined particles were subjected to another round of 3D classification without alignment and with masking of the detergent. The particles from the best 3D class were subjected to several iterative cycles of 3D refinement, CTF refinement and particle polishing, yielding a final post-processed density at 3.55 Å resolution. Local resolution maps were calculated by ResMap implemented in Relion 3.1 (*7, 10*). The detailed steps of cryo-EM processing is depicted in Figure S6.

The data processing of AC8-CaM-Gαs in nanodisc was performed in Relion 3.1.3. All movie stacks were motion corrected using MotionCorr 1.1.0 (*8*) and CTF corrected using Gctf(*9*). The rest of image processing was similar to that of AC8-CaM-Gαs sample in detergent. In brief, a total of 6,127,395 particles were autopicked from three datasets using 3D structured obtained in detergent conditions. After multiple rounds of 2D and 3D clarifications, a set of 33,1737 particles was selected for masked 3D refinement (using masks including or excluding the nanodisc density), resulting in a 3D reconstruction at 3.97 Å resolution. The refined particles were subjected to several iterative cycles of 3D nonalignment classification, 3D refinement, CTF refinement and particle polishing, yielding a small improvement in quality of map without changes in resolution. Local resolution maps were calculated by ResMap implemented in Relion 4.0 (*10, 11*). The detailed steps of cryo-EM processing are depicted in Figure S8.

***Limited proteolysis-mass spectrometry (LiP-MS)***

*Membrane suspension preparation:* A pellet of 2 L induced HEK 293 F GnTI- cells was thawed on ice and resuspended in 50 mL 100 mM HEPES-KOH (pH 7.4), 150 mM KCl, 1 mM $MgCl_2$ (= LiP buffer) with 2 Roche cOmplete protease inhibitor cocktail tablets and DNAse A at a final concentration of 10 µg/mL. The suspension was homogenized using a dounce homogenizer (20 strokes). The sample was further centrifuged at 1,000 x g for 10 min at 4 °C, the supernatant was divided in half, filled up to 50 mL and centrifuged at 35,000 rpm (142,000 x g; Beckman Ti45 rotor) for 40 min at 4 °C. Both pellets were resuspended in a total of 15 mL LiP buffer and homogenized with a dounce homogenizer (20 strokes). The sample was aliquoted, snap frozen in liquid nitrogen and stored at -80 °C until used.

The protein concentration of the membrane suspensions was determined using the BCA assay (Pierce BCA Protein Assay Kit).

*LiP-MS Gβγ titration:* The previously prepared membrane suspensions were thawed on ice and diluted to 2 µg/µL in LiP buffer with 1 mM $CaCl_2$, 1 mM $MnCl_2$ and 100 µM GTPγS. Purified Gβγ was diluted in LiP buffer with 1 mM CaCl2, 1 mM $MnCl_2$ and 100 µM GTPγS and 0.1 % digitonin. All experiments were conducted in quadruplicates. 2 µL of diluted Gβγ at different concentrations (1.5 µg/µL, 1 µg/µL, 0.5 µg/µL, 0.25 µg/µL, 0.05 µg/µL and 0.005 µg/µL) and a buffer blank was spiked into 50 µL of AC8 membrane suspension and incubated for 10 min at 25 °C in a Thermocycler. 5 µL of proteinase K from *Tritirachium album* (Sigma Aldrich) at a concentration of 0.2 µg/µL were added to the samples and incubated for 5 min at 25 °C. Proteinase K was inactivated by heating the samples to 99 °C for 5 minutes, cooling them at 4 °C for 5 min and further addition of 5 % sodium deoxycholate (final concentration).

Additionally, tryptic controls (TC) were produced for the buffer blank treated and the samples treated with the highest amount of Gβγ in quadruplicates: to 50 µL of membrane suspension 2 µL of buffer blank or 2 µL of Gβγ (1.5 µg/µL) were spiked and incubated for 10 minutes. 5 µL of water (instead of proteinase K) were added incubated for 5 min at 25 °C. The samples were then treated the same way as the LiP samples.

*LiP-MS Gαs titration:* The experiment was conducted in a similar way to that with the Gβγ. Briefly, samples were diluted in LiP buffer with 1 mM $CaCl_2$, 1 mM $MnCl_2$ and 100 uM GTPγS and Gαs was titrated in a range of 0-3 µg per sample in the same steps as used for the Gβγ titration.

*Tryptic digest:* All LiP and TC samples were subjected to a tryptic digest. Briefly, disulfide bonds were reduced with 5 mM tris(2-carboxyethyl)phosphine-hydrochloride for 40 minutes at 37 °C with slight agitation. Reduced cysteines were alkylated with 40 mM iodoacetamide at room temperature in the dark with slight agitation. The samples were diluted to a final sodium deoxycholate concentration of 1% with 100 mM ammonium bicarbonate. 1 µL of Lysyl endopeptidase LysC (1 µg/µL) and 2 µL of sequencing grade trypsin (0.5 µg/µL) were added and the samples were incubated overnight at 37 °C with slight agitation. The next day, the digest was stopped by adding formic acid to a final concentration of 2%. The precipitated sodium deoxycholate was removed by filtering through a 96-well Corning® 2 µM PVDF plate and the samples were desalted on a 96-well MacroSpin plate (The Nest Group). The peptides were eluted with 80 % acetonitrile, 0.1 % formic acid and dried in a vacuum centrifuge. The samples were stored at -20 °C until further use.

LC-MS/MS: The samples were reconstituted in 5% ACN, 0.1% FA with the addition of iRT peptides (Biognosys). For library generation, pooled samples of all replicates of one condition were measured in DDA (data dependent acquisition). The unpooled samples were measured in DIA (data independent acquisition). Gβγ titration data was acquired on an Orbitrap Fusion Tribrid mass spectrometer (ThermoFisher Scientific), equipped with a nanoelectrospray source and an Easy-nLC 1200 nanoflow LC system (Thermo Scientific). Peptides were separated on a 40 cm x 0.75 i.d. column packed in-house with 1.9 um C18 beads (Dr. Maisch Reprosil-Pur 120) using a linear gradient from 3-35% B (A: 0.1% FA, B: 95% ACN, 0.1% FA) over 120 min at 300 nL/min. The column was heated to 50 °C. DIA samples were acquired with a 41 window (1 m/z overlap) method. For DDA measurements the Orbitrap resolution was set to 120,000, the scan range was between 350 and 2000 m/z. Maximum injection time was 50 ms, the normalized AGC target was set to 200%, RF lens was set to 60%. Exclusion time was 30s, charge states between +2 and +6 were analyzed. The fragmentation method was higher energy collisional dissociation (HCD) with a collision energy of 30 % and a scan range of 150-2000 m/z.For DIA measurements a survey MS1 scan was acquired at an Orbitrap resolution of 120,000 with a scan range between 350 and 1400 m/z and a maximum injection time of 100 ms. The normalized AGC target was at 200 %, RF lens at 30 %. Consecutively, 41 variably sized MS2 windows with 1 m/z overlap were acquired. The Orbitrap resolution was set to 30,000, the HCD collision energy to 30 %. The normalized AGC target was set to 100%.

The Gαs titration samples were acquired on an Orbitrap Eclipse Tribrid mass spectrometer (ThermoFisher Scientific). For DDA measurements the Orbitrap resolution was set to 120,000, the scan range was between 350 and 1400 m/z. Maximum injection time was 54 s, the normalized AGC target was set to 200%. RF lens was set to 30% and charge states between +2 and +7 were analyzed. The exclusion time was set to 20 s. The fragmentation method was HCD with a collision energy of 30 %. The Orbitrap resolution was set to 30,000, the scan range was between 150 and 2000 m/z. Maximum injection time was 54 ms, the normalized AGC target was set to 200 %. For DIA measurements a survey MS1 scan was acquired with the Orbitrap resolution set to 120,000 and a scan range between 350 and 1400 m/z. The injection time was set to 100 ms, the normalized AGC target to 200 %, RF lens was at 30 %. The DIA window setup was the same as mentioned above for the Orbitrap Fusion Tribrid mass spectrometer method.

*Data analysis:* The acquired raw data was analyzed using Spectronaut v. 15.4(12). Tryptic control data and LiP data was searched separately. A library was produced from searches of DDA data. Default search parameters were slightly altered: a minimum peptide length of 5 amino acids was applied and single hits were excluded, missing values were not imputed. For LiP-MS data, protease specificity was set to "semi-specific". Data was exported from Spectronaut and further analyzed in R. Quality control, data analysis and interpretation was conducted using the R packages protti (13), tidyverse (14), data.table (15). The tryptic control data was checked for AC8 protein abundance changes and four-parameter dose response curves were fit onto all AC8 peptides detected in the LiP samples using an implementation of the drc (16) R package. Pearson correlation was calculated and peptides with a Pearson correlation r of >0.85 were considered significant. Peptides for which only the one concentration point was different to the rest were not considered significant.

#### *Crosslinking-mass spectrometry (XL-MS)*

AC8, G$\alpha$s and G$\beta\gamma$ were concentrated and the purification buffer was exchanged for 50 mM HEPES pH 7.5, 150 mM NaCl, 0.02 % GDN, 0.5 mM $CaCl_2$, 1 mM $MgCl_2$, 2.5 mM $MnCl_2$, 100 uM GTP$\gamma$S. AC8 crosslinking was performed with 60 µg protein, for the other conditions, 30 µg AC8 was used. G proteins were incubated with 100 uM GTP$\gamma$S for 10 min at 25 °C prior to crosslinking. The interactor reactions were conducted at molar rations of AC8: G$\alpha$s 1:1.5, AC8:G$\beta\gamma$ 1:3, AC8:CaM 1:2, AC8:CaM: G$\alpha$s 1:2:1.5. All samples were incubated for 30 min at 25 °C.

Primary amine crosslinking was performed with 1 mM DSS (disuccinimidyl suberate) (11.4 Å linker length (*17*))at a 1:1 ratio of DSS-$d_0$ and DSS-$d_{12}$ (Creative Molecules; 25 mM stock in dimethyl formamide). Samples were incubated at 25 °C for 1 hour at 300 rpm. The crosslinking reaction was quenched with 50 mM ammonium bicarbonate.

Carboxyl group and primary amino group crosslinking was performed with 44 mM PDH (pimelic dihydrazyde)-$d_0$ (ABCR) mixed at a 1:1 ratio with PDH-$d_8$ (Sigma-Aldich) (12.3 Å linker length (*18*)) and DMTMM (4-(4,6-dimethoxy-1,3,5-triazin-2-yl)-4-methylmoorpholinium chloride. Samples were incubated at 25 °C for 1 hour at 300 rpm. To quench the reaction, the samples were desalted using Zeba Spin Desalting columns. An aliquot of each crosslinked and non-crosslinked samples was analyzed with SDS-PAGE (Fig. S11).

All crosslinked samples were dried in a vacuum centrifuge before further processing. The dried samples were resuspended in 50 µL 8 M urea, disulfide bonds were reduced with 2.5 mM tris(2-carboxyethyl)phosphine hydrochloride for 30 min at 37 °C. Free thiol groups were alkylated with 5 mM iodoacetamide for 30 min at room temperature in the dark. 25 µL of 150 mM ammonium bicarbonate were added to the samples, followed by the addition of 0.6 µg LysC (1:100 enzyme:protein). After 2 hours incubation at 37 °C 320 µL of 50 mM ammonium bicarbonate were added to the samples. Trypsin was added at an enzyme to protein ratio of 1:50. Samples were incubated overnight at 37 °C with slight agitation. Formic acid was added to a concentration of 2 % and the samples were desalted with solid-phase extraction (MicroSpin Columns 5-60 µg capacity). The samples were evaporated to dryness and resuspended in 30 % ACN, 0.1 % trifluoroacetic acid (TFA).

The samples were further fractionated using size-exclusion chromatography (Superdex 30 Increase column 300 x 3.2 mm, GE Healthcare) in 30 % ACN, 0.1 % TFA at a flow rate of 50 µL/min. Four fractions were collected for each sample and dried in a vacuum centrifuge.

*LC-MS/MS:* 10 % of each SEC fraction – except for the highest molecular weight fractions of AC8-CaM (PDH/DMTMM) and AC8-CaM-Gαs (PDH/DMTMM) (2.5 %) – were injected onto an Easy nLC-1200 HPLC (Thermo Scientific) system coupled to an Orbitrap Fusion Lumos mass spectrometer (Thermo Scientific). Peptides were separated on an Acclaim PepMap RSLC C18 column (250 mm x 75 um; Thermo Scientific) with a gradient from 11-40% B (A: 2 % ACN, 0.15 % FA; B: 80 % ACN, 0.15 % FA) in 60 min, at a flow rate of 300 nL/min.

All samples were acquired in DDA with a cycle time of 3 seconds and a dynamic exclusion of 30 seconds. MS1 Orbitrap resolution was set to 120,000, charge states between +3 and +7 were selected in a range of 350-1500 m/z and fragmented with collision-induced dissociation in a linear ion trap at a collision energy of 35 %. Fragment ions were detected in in the ion trap.

In addition, PDH/DMTMM samples were measured in duplicates with a method similar to the method above but with fragment detection in the Orbitrap at a resolution of 30,000.

*Data analysis:* To determine which proteins were present in the sample, the smallest molecular weight fractions of DSS crosslinked samples from each condition were searched with Spectromine v.3 (Biognosys) against a database of human, bovine, *Trichoplusia ni* and target proteins, as well as common contaminants (used from MaxQuant(*19*)). Identified proteins with > 5% LFQ intensity (compared to target proteins) were selected for the initial FASTA files. The initial FASTA files were used for crosslink identification with xQuest v. 2.1.5 (*20*). The databases were reversed and shuffled. Trypsin was set as the digestion enzyme, maximum number of missed cleavages 2, carbamidomethylation on cysteines was set as a fixed modification, oxidation of methionines was set as a variable modification, peptide lengths were set to 4-40 amino acids. MS1 error tolerance was set to 15 ppm, MS2 tolerance was set to 0.2 Da. All proteins with identified monolinks were included in a FASTA file for the main crosslinking search.

For the main searches settings similar to the initial search were applied with trypsin as the digestion enzyme, an MS1 error tolerance of ± 15 ppm, and an MS2 error tolerance of ± 0.2 Da. Crosslinked amino acids for DSS crosslinking were lysines and the protein N-terminus, for PDH aspartate and glutamate and for DMTMM lysines with aspartate and lysine with glutamate. The error tolerances were changed for the measurements that employed orbitrap detection. MS2 tolerance, crosslink MS2 tolerance, and peak matching tolerances (cp_tolerance, cp_tolerancexl) were set to ± 15 ppm. The minimum peak number was set to 10.

Crosslinked peptides were filtered with a mass tolerance window of ± 5 ppm, TIC ≥0.1, minimally 3 matched ions and a delta score of <0.9. To achieve <5 % false discovery rate, the xQuest score cutoff was set at less than 5 % of decoy hits compared to target hits. All spectra were checked and filtered manually with at least three bond cleavages. For AC8 heteromeric/inter-protein XLs, homomeric/intra-protein XLs, monolinks and homomultimeric links/intermolecular selflinks were exported. For the interactors we only exported hetero- and homomeric XLs, as well as homomultimeric links (dimerization links). Crosslinks between 2 peptides sharing the same sequence were considered to be intermolecular selflinks.

*Data visualization:* All DSS, PDH and DMTMM crosslinks were combined in one file and used for visualization in pyMOL v. 2.4.1 using the plugin PyXlinkViewer (*21*). 30 Å were set as violation cutoff. This cutoff was defined based on the maximal Cα-Cα distance reported for DSS (*17*). Distances calculated by PyXlinkViewer were exported and used for visualization with xiNET (*22*).

#### *AC8-Gβγ docking*

For docking studies, we used the Gβγ structure determined by Davis et al. (*23*) (PDB: 1xhm). All three crosslinks between structured regions of AC8 and Gβγ were considered for docking.

HADDOCK 2.4(*24*) was used to predict the AC8-Gβγ complex using the detected inter-protein crosslinks as distance restraints, keeping the default settings for protein-protein docking. DisVis (*25*) was used to visualize the accessible interaction space, check for false positives and to further calculate the residues involved in the interaction. The distances set in the restraints file were 8.7-20.5 Å for PDH and 6.5-16 Å for DMTMM (*18*). The residues used for docking were residue 674 and 678 on AC8 (PDH) and AC8 residue 936 (DMTMM). All 3 crosslinked with residue 312 on the Gβ subunit. GetArea (*26*) was used to calculate the per residue surface accessibility of AC8 and Gβγ and residues with at least 40 % accessibility were considered for the analysis.

Cluster 1 showed the lowest mean van der Waals energy, the highest fraction of common contacts and a plausible orientation of Gβγ that would facilitate membrane anchoring (**Fig. S15**). That is why cluster 1, structure 1 was used for illustration purposes.

#### *AlphaFold-Multimer prediction*

An AlphaFold-Multimer (AlphaFold v. 2.2.0) prediction of AC8 (UniProt ID: E1BQ12) interacting with CaM (UniProt ID: P62157) and Gαs (UniProt ID: P04896) and was run on Euler scientific computing cluster (ETH Zurich) using the standard submission script for SLURM submission and resource calculation (https://gitlab.ethz.ch/sis/alphafold_on_euler).

### SUPPLEMENTARY FIGURES

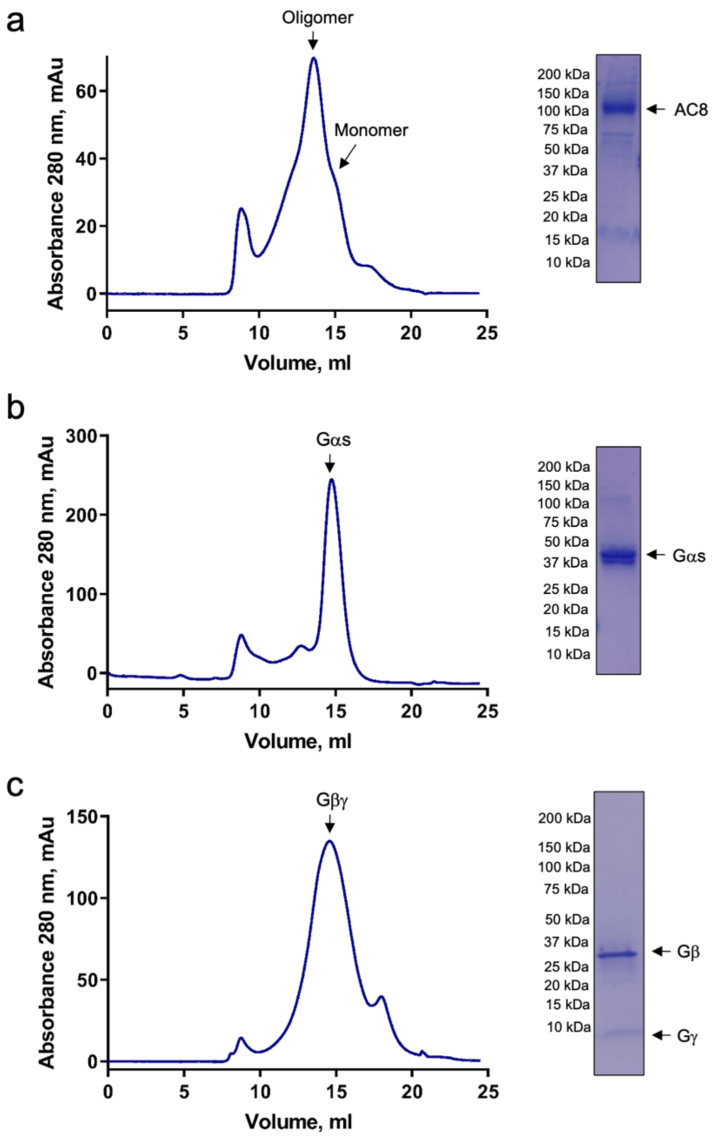


**Figure S1. Recombinant expression and purification of AC8 and G-protein subunits.** Size exclusion chromatography (SEC) and SDS-PAGE analysis of purified **(a)** bovine AC8, **(b)** G protein Gαs subunit, **(c)** G protein Gβγ subunit. SEC for AC8 and G-proteins was performed using a Superose 6 Increase column.

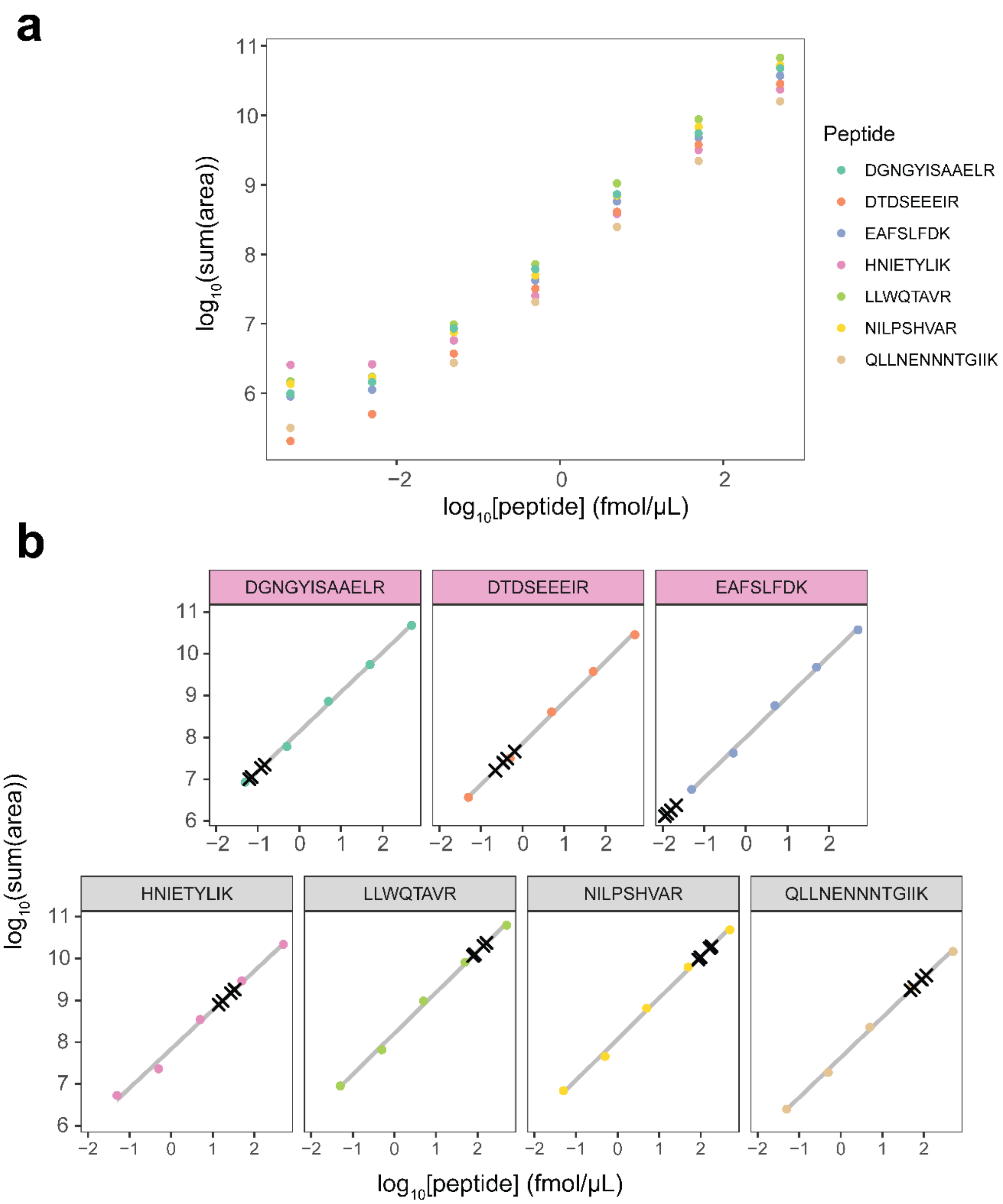


**Figure S2. PRM calibration curves for AC8 and CaM heavy labeled peptides. (a)** All calibration points for AC8 (HNIETYLIK, LLWQTAVR, NILPSHVAR, QLLNENNNTGIIK) and CaM (DGNGYISAAELR, DTDSEEEIR, EAFSLFDK) heavy labeled peptides. **(b)** Calibration curves for CaM peptides (pink header) and AC8 peptides (grey header). Summed up intensities of the 5 highest ranked fragments of the 0.5 µL and 1 µL injections detected in the samples are shown as a black crosses.

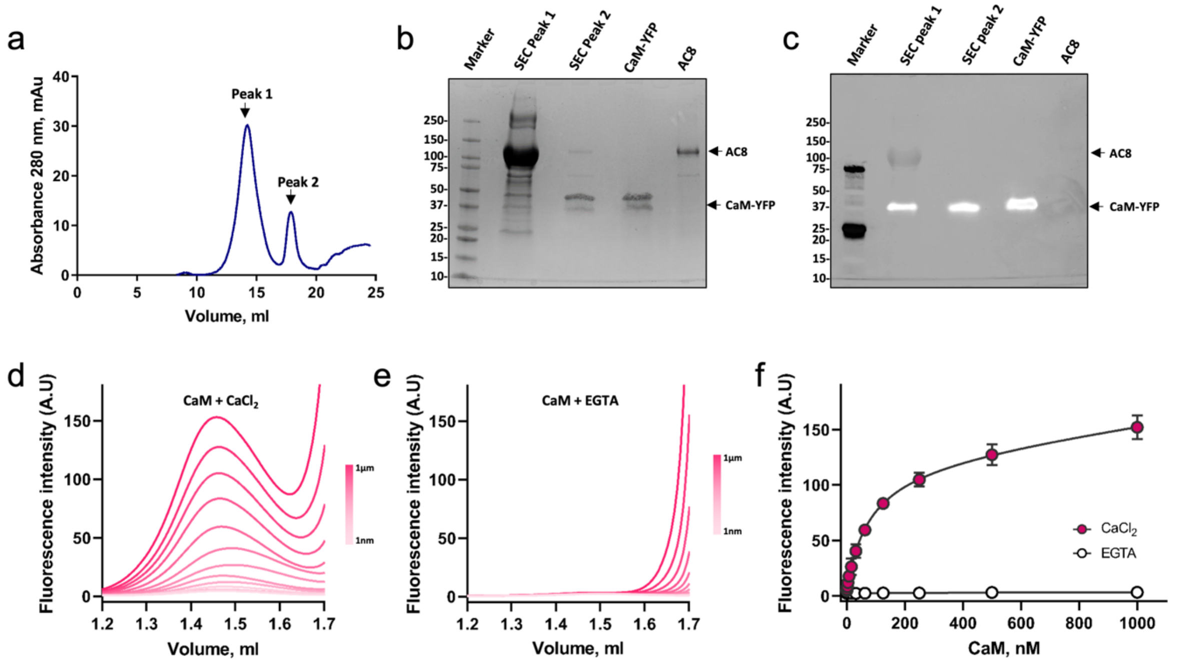


**Figure S3. Purified AC8 binds CaM with high affinity in the presence of $Ca^{2+}$. (a)** The size exclusion chromatography (SEC) profile and **(b-c)** SDS-PAGE analysis of AC8-$Ca^{2+}$/CaM-YFP complex. The complex was prepared by mixing purified recombinant AC8 and CaM-YFP in 1:2 ratio, respectively. The panel b shows Coomassie stained SDS-PAGE of AC8-$Ca^{2+}$/CaM complex after SEC. Panel c shows overlayed prestained and YFP-fluorescence image of SDS-PAGE displayed in panel b. **(d-e)** Representative FSEC profiles showing specific increase in fluorescence intensity in AC8 elution peak with increasing concentration of CaM. **(f)** The saturation-binding curves shows nanomolar affinity (77.7 ± 10.9 nM, n = 3) of AC8 for CaM in presence of calcium and lack of AC8-CaM interaction in absence of calcium. For the experiments in panel f, the data are shown as mean ± S.E.M (n = 3).

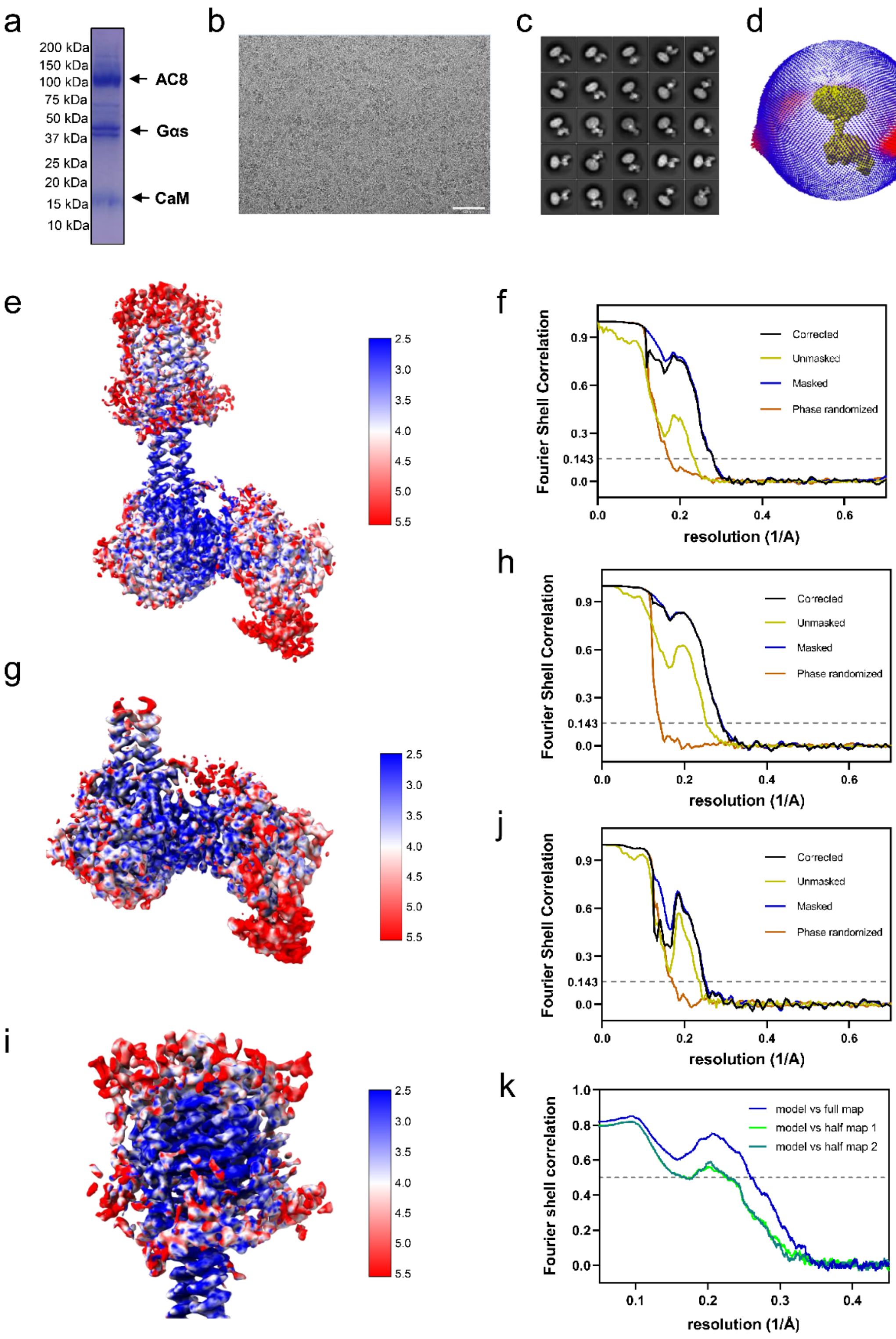


**Figure S4. Cryo-EM data analysis of AC8-$Ca^{2+}$/CaM-Gαs-Forskolin-MANT-GTP in detergent. (a)** SDS-PAGE analysis of AC8-$Ca^{2+}$/CaM-Gαs (AC8-CaM-Gαs) complex used for cryo-EM analysis. **(b)** A representative micrograph of AC8-CaM-Gαs complex in the presence of 0.5 mM Forskolin, 0.5 mM MANT-GTP, 5 mM $MnCl_2$, 2 mM $MgCl_2$ and1 mM $CaCl_2$. **(c)** Representative 2D classes of AC8-CaM-Gαs. **(d)** Angular distribution histogram of AC8-CaM-Gαs complex dataset. **(e-f)** Local resolution and

FSC curve for AC8-CaM-G$\alpha$s complex after subtraction of detergent micelle. **(g-h)** Local resolution and FSC curve of the soluble domain of AC8 and G$\alpha$s after focus refinement without TM domain and detergent micelle. **(i-j)** Local resolution and FSC curve for TM domain after focus refinement without detergent micelle. **(k)** Map to model FSC plot of soluble domain of AC8-bound to G$\alpha$s.

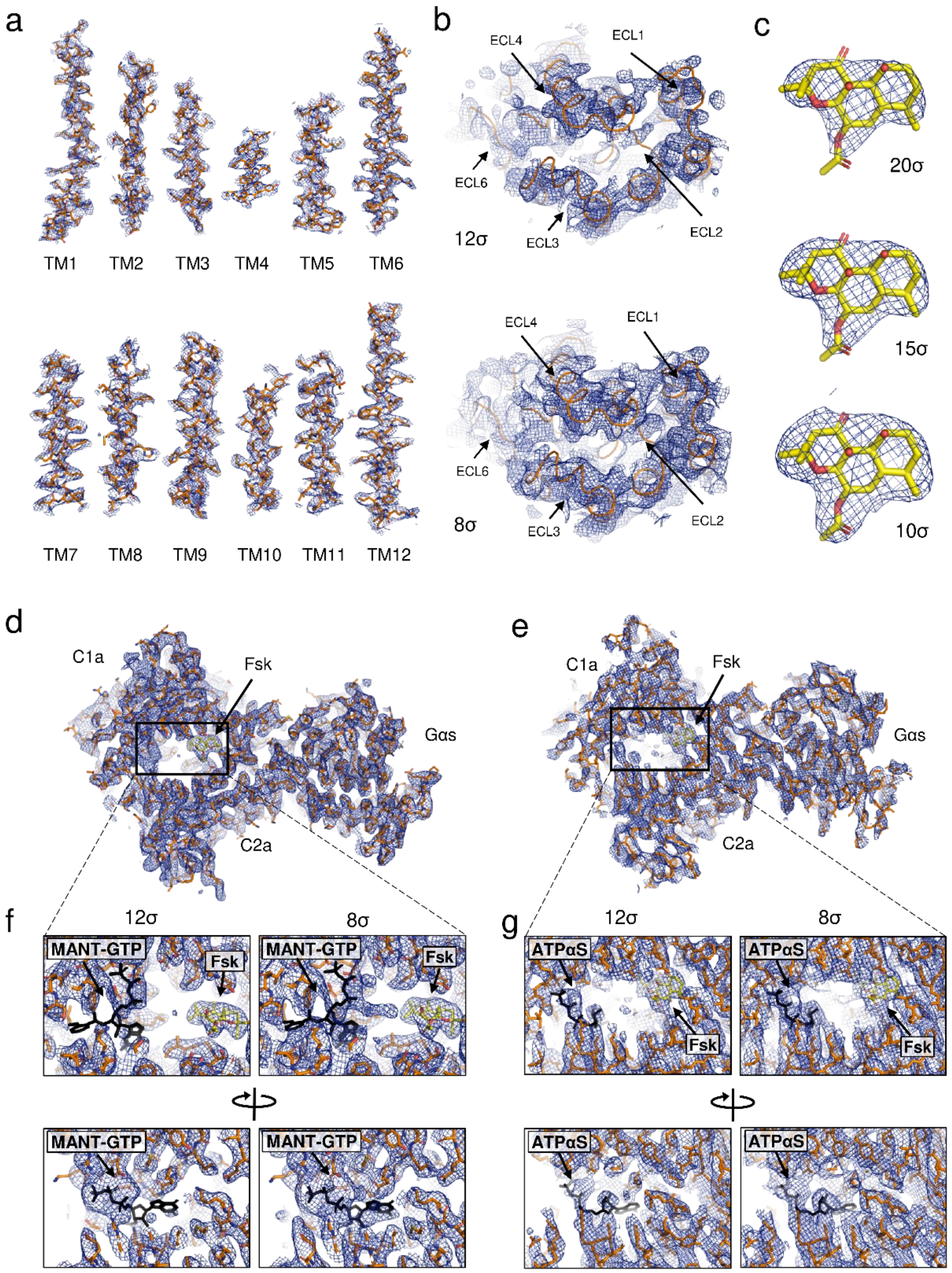


**Figure S5. Cryo-EM map features of bovine AC8 bound to CaM and Gαs. (a)** Isolated density map for 12 TM helices of AC8-CaM-Gαs complex in GDN micelles contoured at 10σ threshold level. **(b)** Cryo-EM density features of extracellular loops contoured at 12σ and 8σ levels. **(c)** Cryo-EM density features of forskolin contoured at 20σ, 15σ and 10σ levels. **(d-e)** Cryo-EM density features for the catalytic domain of AC8 bound to Gαs in GDN micelle (panel d) and MSP1ED1 nanodisc (panel e). **(f-g)** Depiction of unresolved density features of MANT-GTP (panel f) and ATPαS (panel g) contoured at 12σ and 8σ levels. MANT-GTP and ATPαS binding poses (shown as black sticks) are, based on structures of chimeric AC5C1a-AC2C2a (PDB: 3C16, 3MAA)

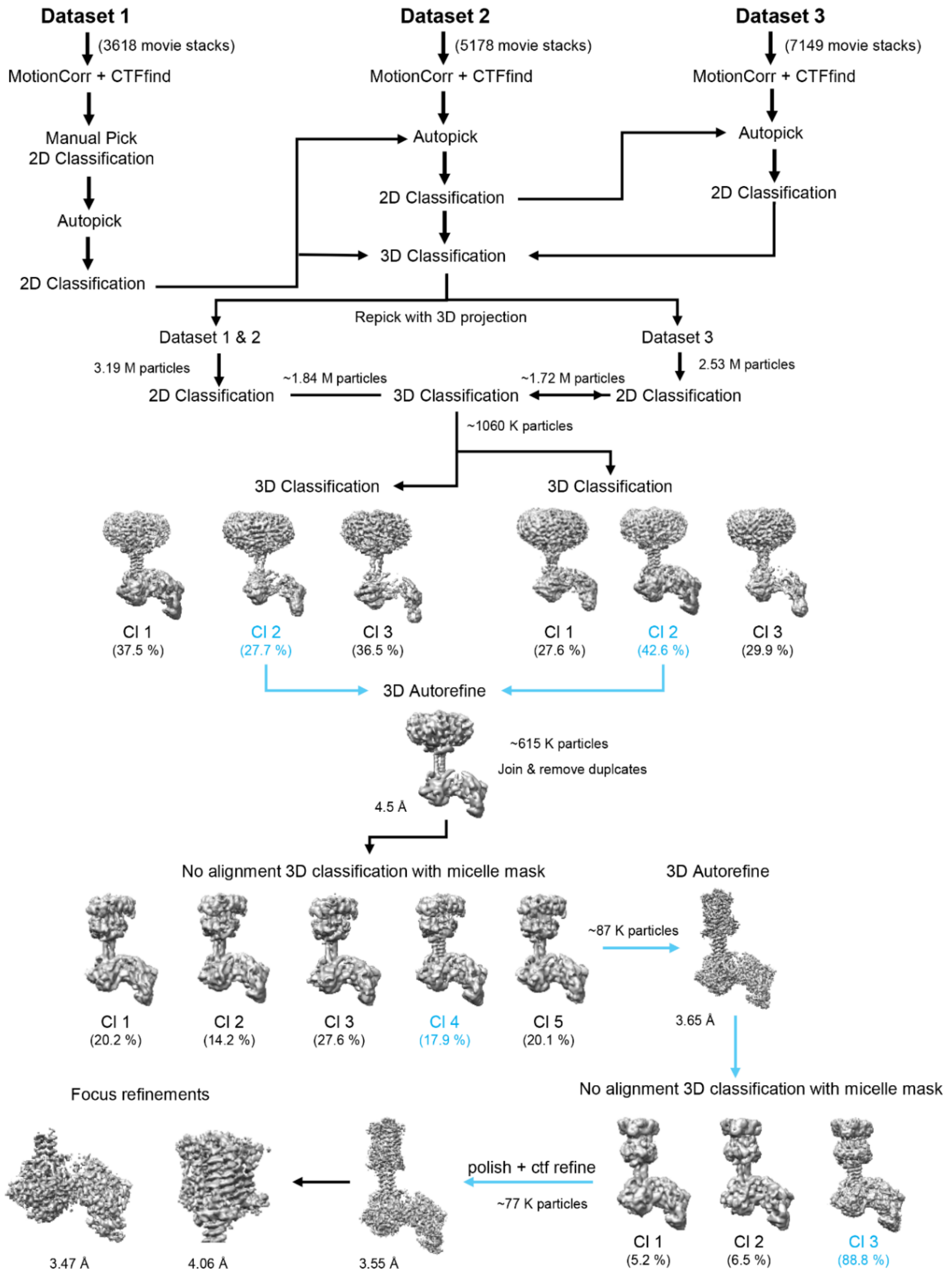


**Figure S6. Cryo-EM data processing pipeline of AC8-$Ca^{2+}$/CaM-G$\alpha$s-Forskolin-MANT-GTP complex in the detergent micelle.**

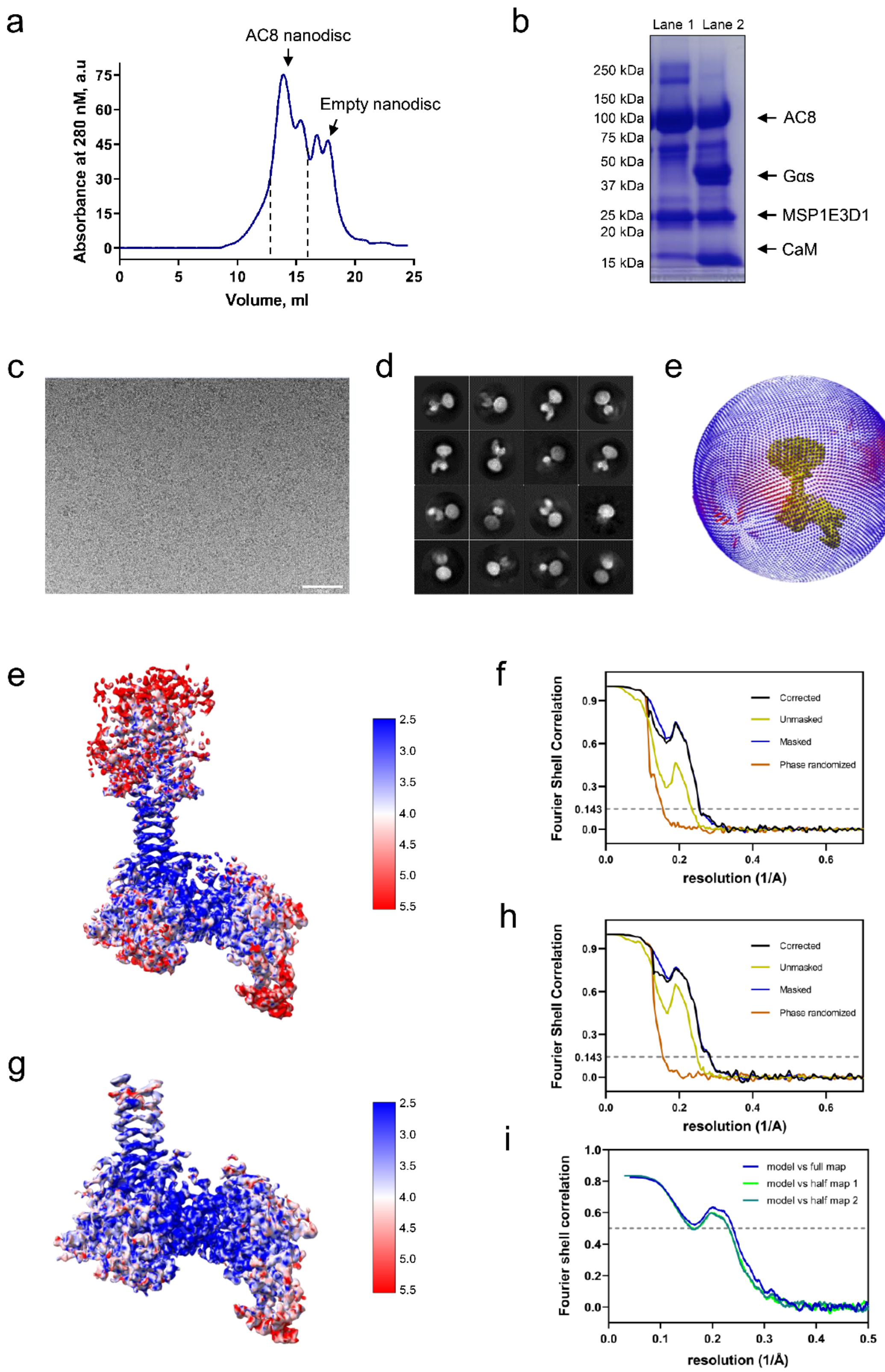


**Figure S7. Cryo-EM data analysis of AC8-$Ca^{2+}$/CaM-Gαs-Forskolin-ATPαS complex in lipid nanodisc. (a)** Size exclusion chromatography (SEC) of AC8 reconstituted in lipid nanodisc with brain polar lipid and MSP1E3D1 **(b)** SDS-PAGE analysis of AC8 nanodisc (lane 1) and cryo-EM sample of AC8-$Ca^{2+}$/CaM-G$\alpha$s complex in lipid nanodisc (lane 2). **(c)** A representative micrograph of AC8-CaM-G$\alpha$s complex in the presence of 0.5 mM Forskolin, 1 mM ATP$\alpha$S, 5 mM $MnCl_2$, 2 mM $MgCl_2$ and 1 mM $CaCl_2$. **(d)** Representative 2D classes of AC8-CaM-G$\alpha$s in nanodisc. **(e)** Angular distribution histogram of AC8-CaM-G$\alpha$s complex in nanodisc. (**f-g)** Local resolution and FSC curve for AC8-CaM-G$\alpha$s complex after subtraction of detergent micelle. **(h-j)** Local resolution, FSC curve of the soluble domain of AC8 and G$\alpha$s after focus refinement without TM domain and detergent micelle and map to model FSC plot of soluble domain of AC8-bound to G$\alpha$s.

.

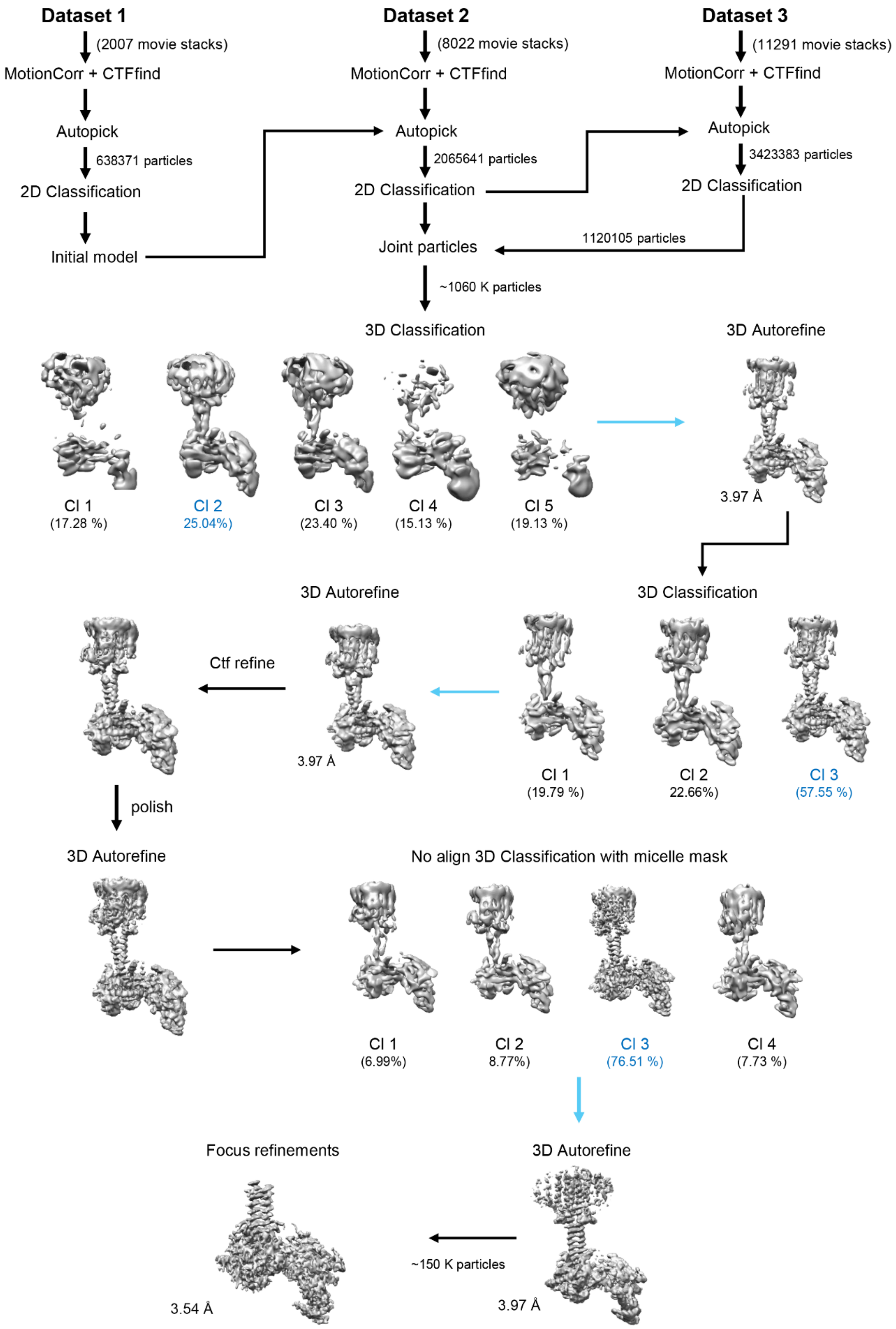


**Figure S8. Cryo-EM data processing pipeline of AC8-$Ca^{2+}$/CaM-Gαs-Forskolin-ATPαS complex in lipid nanodisc.**

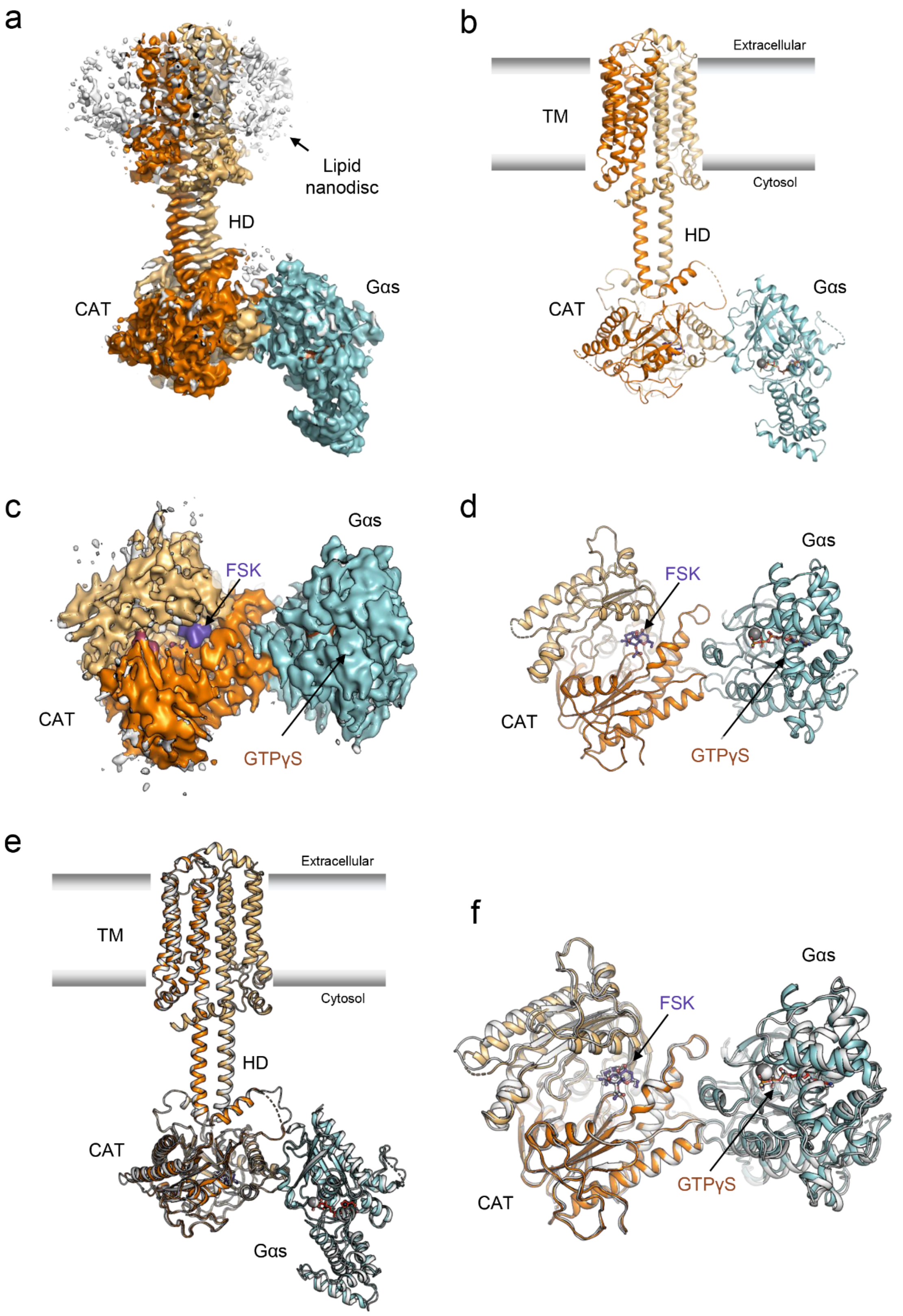


**Figure S9. Cryo-EM structure of AC8-CaM-Gαs structure in lipid nanodisc. (a-d)** Cryo-EM map and model of AC8-CaM-Gαs-Forskolin-ATPαS complex reconstituted in lipid nanodisc (MSP1E3D1 and brain polar lipid). (**e-f**) Comparison of the cryo-EM structures of AC8-Gαs complex resolved in detergent (white) and lipid nanodisc (orange).

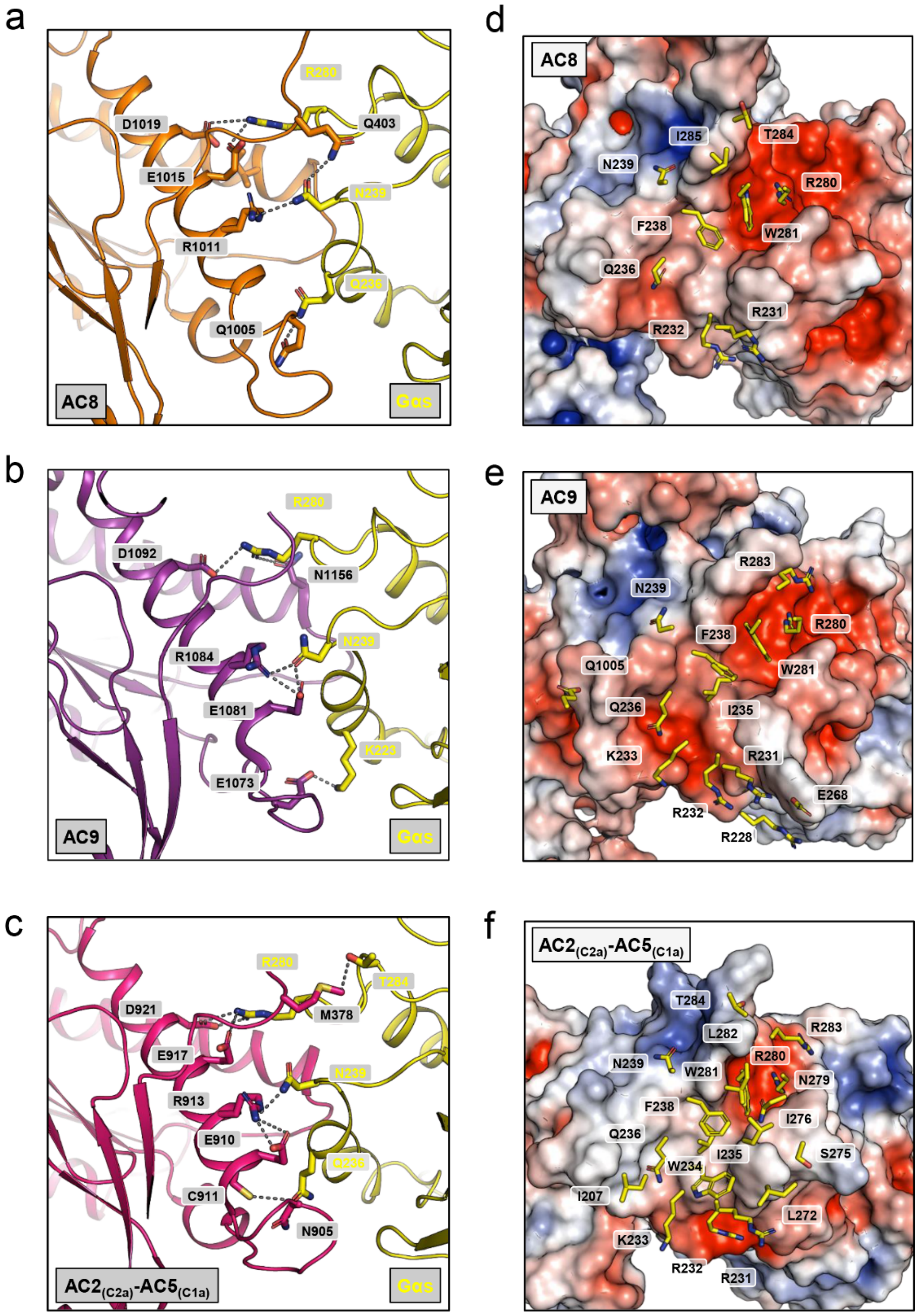


**Figure S10. Gαs binding interfaces in ACs**. **(a-c)** Depiction of AC-Gαs interfacial residues observed in the cryo-EM structure of AC8-CaM-Gαs (top), AC9-Gαs (middle), and crystal structure of chimeric $AC2_{(C2a)}$-$AC5_{(C1a)}$-Gαs complex (bottom). (**d-f)** Electrostatic surface representation of AC8 (top), AC9 (middle), and chimeric $AC2_{(C2a)}$-$AC5_{(C1a)}$ (bottom) showing interfacial residues of Gαs (yellow color) within 4 Å of AC.

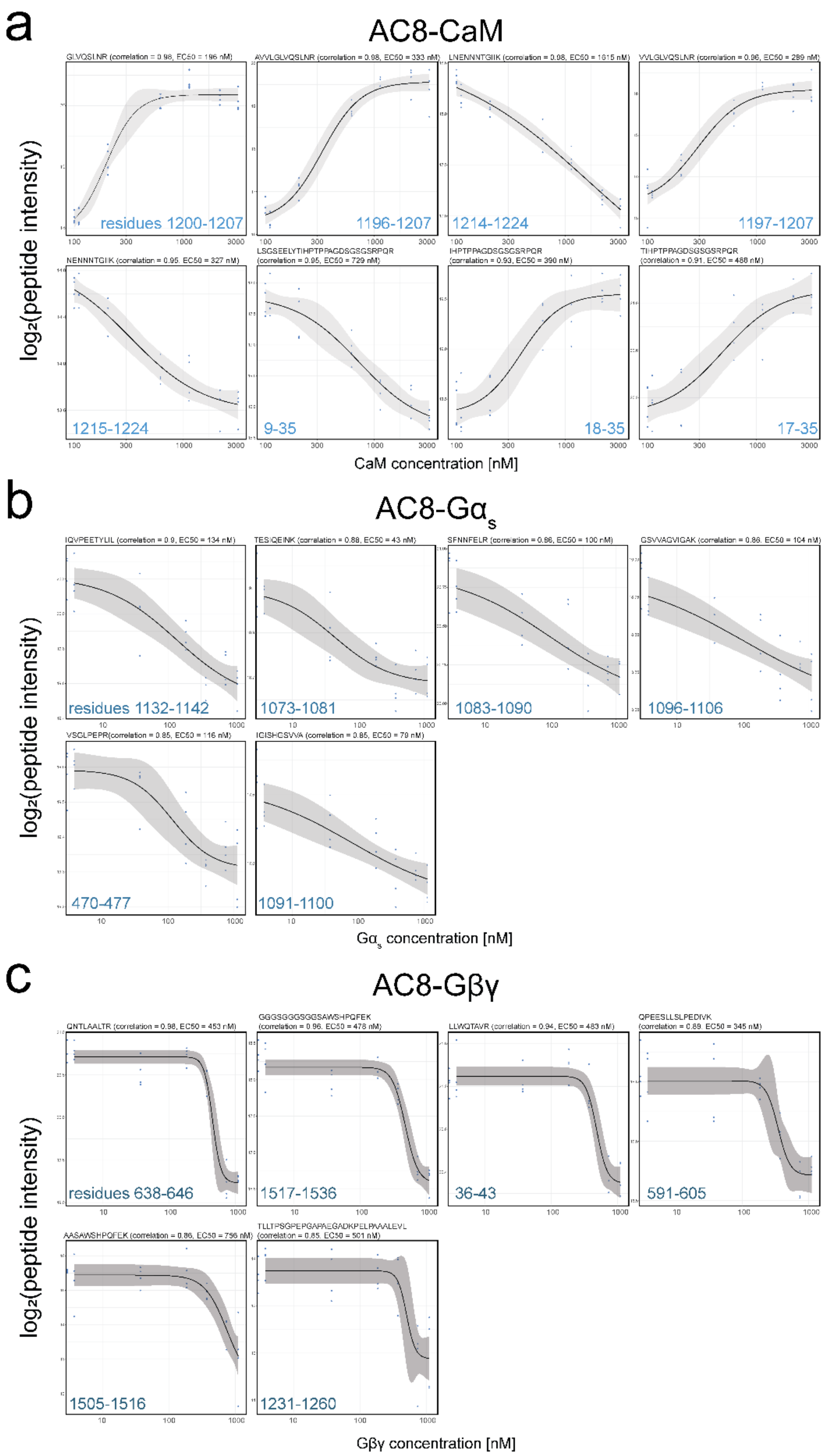


**Figure S11. LiP-MS curves for all peptides with Pearson r > 0.85 for AC8-CaM (a), AC8-Gαs (b) and AC8-Gβγ (c) titrations.** Peptide sequences are shown in the header of each plot, alongside the Pearson r (correlation) and $EC_{50}$ values. The peptide positions within the AC8 sequence are indicated at the bottom of each plot. The measured and log2 transformed peptide intensities for each replicate at each interactor concentration are shown as blue dots. The 95 % model confidence interval is shown as a grey shaded area. Peptides with only one changing condition were excluded.

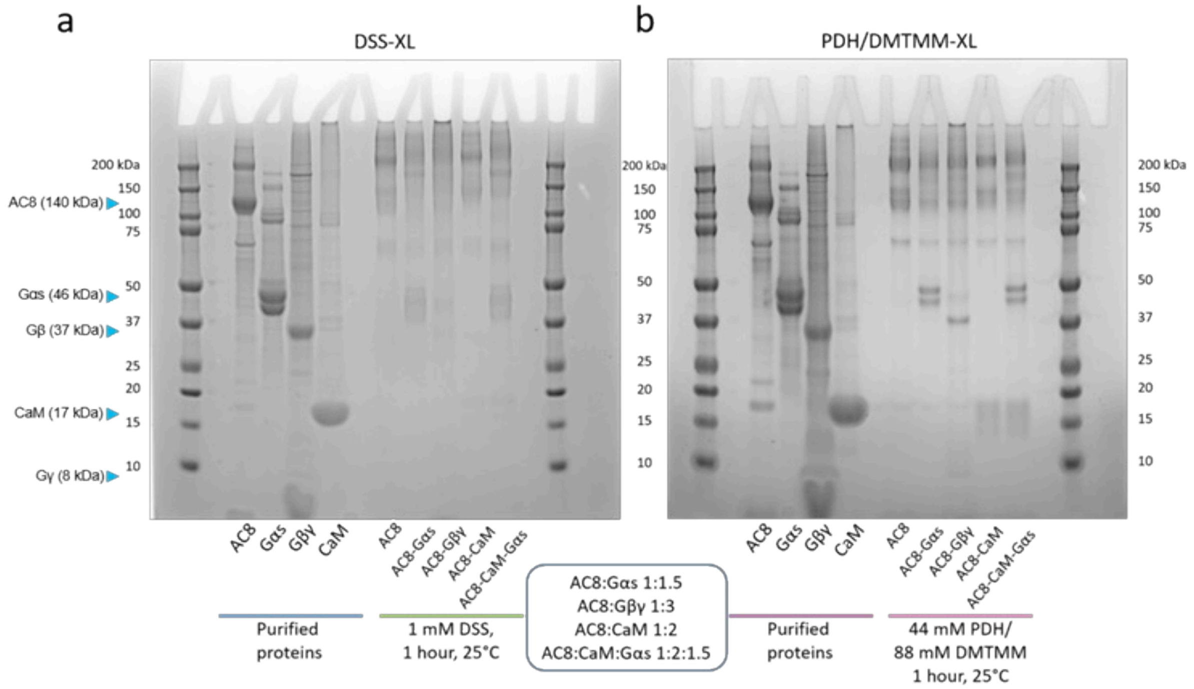


**Figure S12. SDS-PAGE of purified and crosslinked proteins (DSS and PDH/DMTMM) (a)** SDS-PAGE of purified and DSS-crosslinked proteins. The migration distances of the respective proteins are indicated with a blue arrow on the left side of the gel. **(b)** SDS-PAGE of purified and PDH/DMTMM-crosslinked proteins**.** Molar ratios of the crosslinked proteins are listed in the blue box in the center. Reaction conditions are listed below.

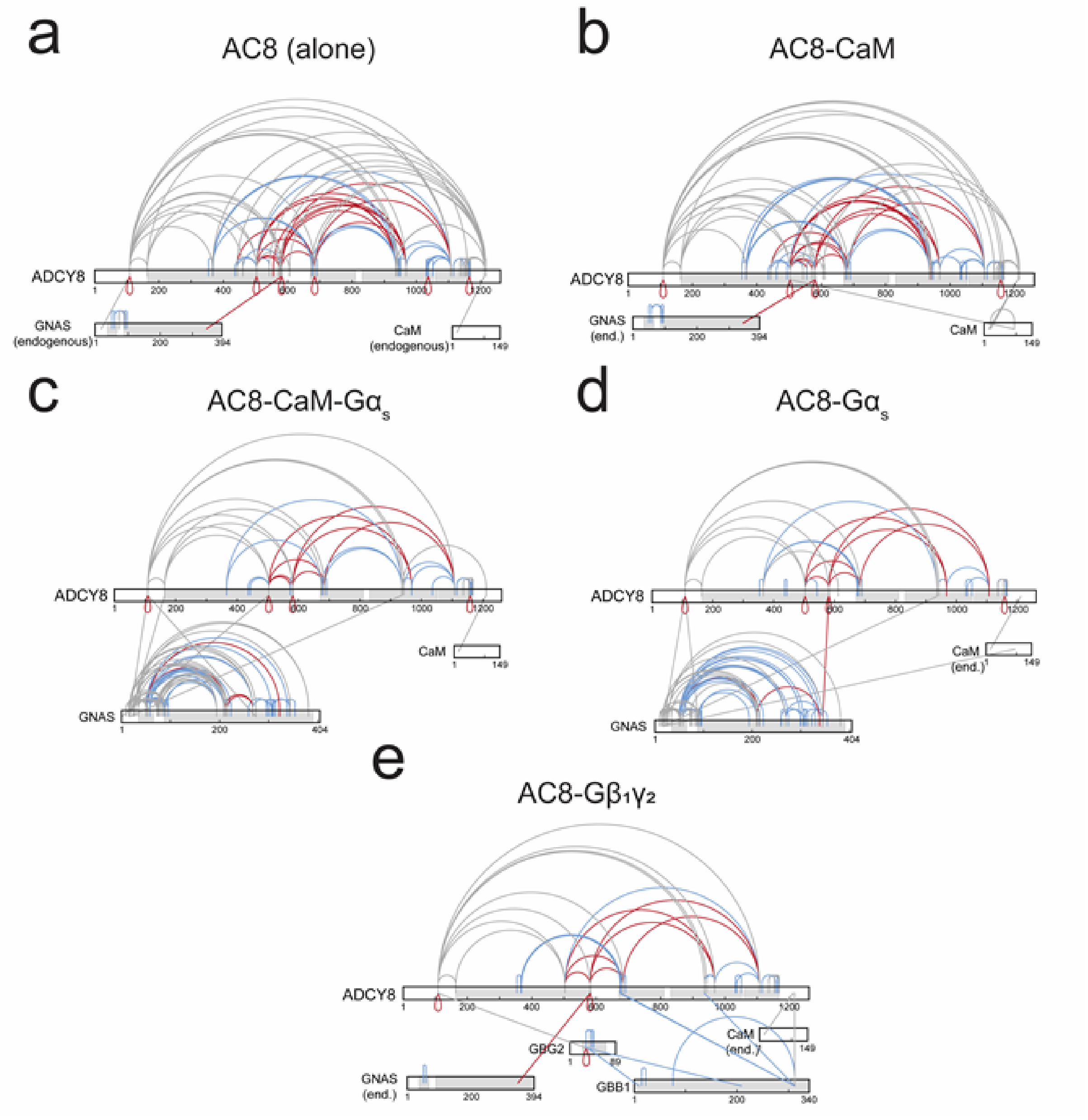


**Figure S13. All detected crosslinks (DSS and PDH/DMTMM). (a-e)** Violated crosslinks (>30Å) are coloured in red, satisfied crosslinks between structured/resolved regions are coloured in blue. Crosslinks that involve flexible regions and regions that are not resolved in the cryo-EM or in the structures used for protein-protein docking are coloured in grey. Homomultimeric links (oligomerization links) are indicated with red drops. Regions with available structure are highlighted in grey on the respective protein sequence. **(a)** crosslinked AC8 with copurified Gαs and CaM. **(b)** crosslinked AC8 and CaM with copurified Gαs **(c)** crosslinked AC8, CaM and Gαs **(d)** crosslinked AC8 and Gαs with copurified CaM **(e)** crosslinked AC8 and Gβγ with copurified Gαs and CaM.

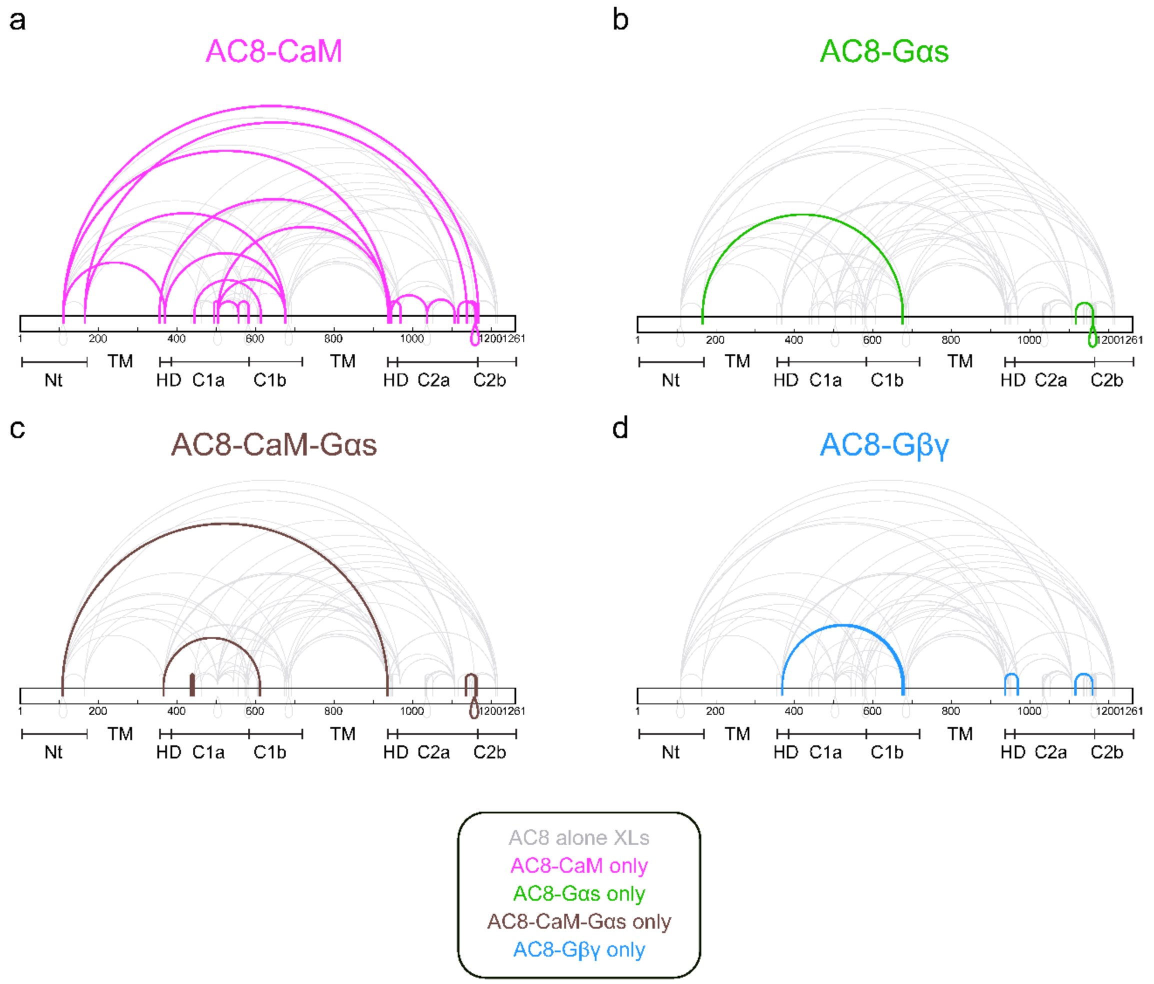


**Figure S14. XL-MS comparison of identified crosslinks between AC8 by itself and interactors. (a)** AC8 alone crosslinks are coloured in light grey, positioned in the background. Crosslinks only found in the AC8-CaM sample (not in the AC8 alone sample) are colored in pink. AC8 domains and their respective locations are indicated below. **(b)** Crosslinks only found in the AC8-G$\alpha$s sample are coloured in green. **(c)** Crosslinks identified only in the AC8-CaM-G$\alpha$s sample are highlighted in brown. **(d)** Crosslinks unique to the AC8-G$\beta\gamma$ sample are coloured in blue.

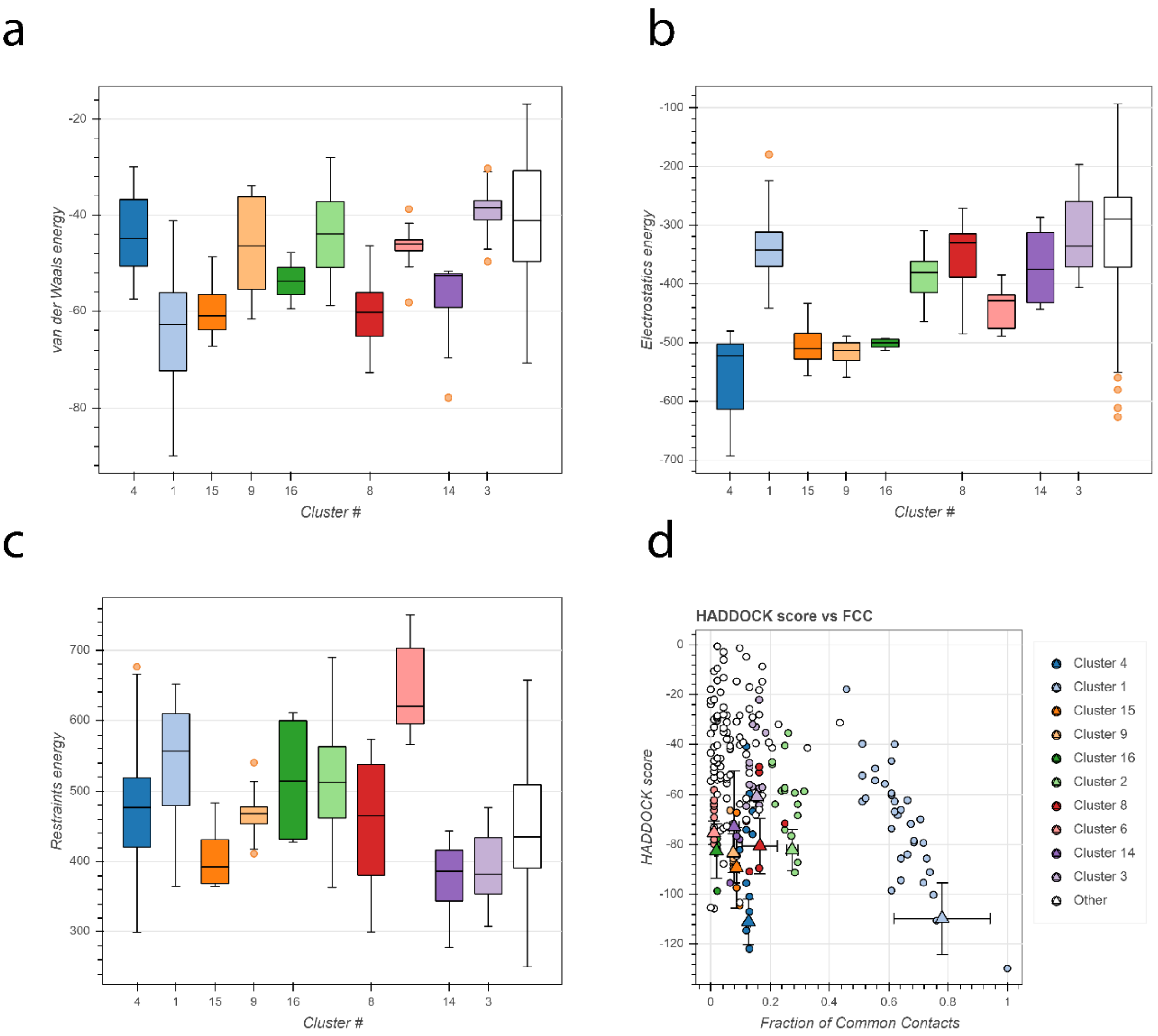


**Figure S15. HADDOCK quality control plots.** The plots were generated by HADDOCK 2.4 directly. **(a)** Van der Waals energy ranges for the different docking clusters generated by HADDOCK. **(b)** Electrostatics energy distributions for the different clusters. **(c)** Restraints energies for the generated clusters **(d)** Dotplot showing the HADDOCK score vs. fraction of common contacts.

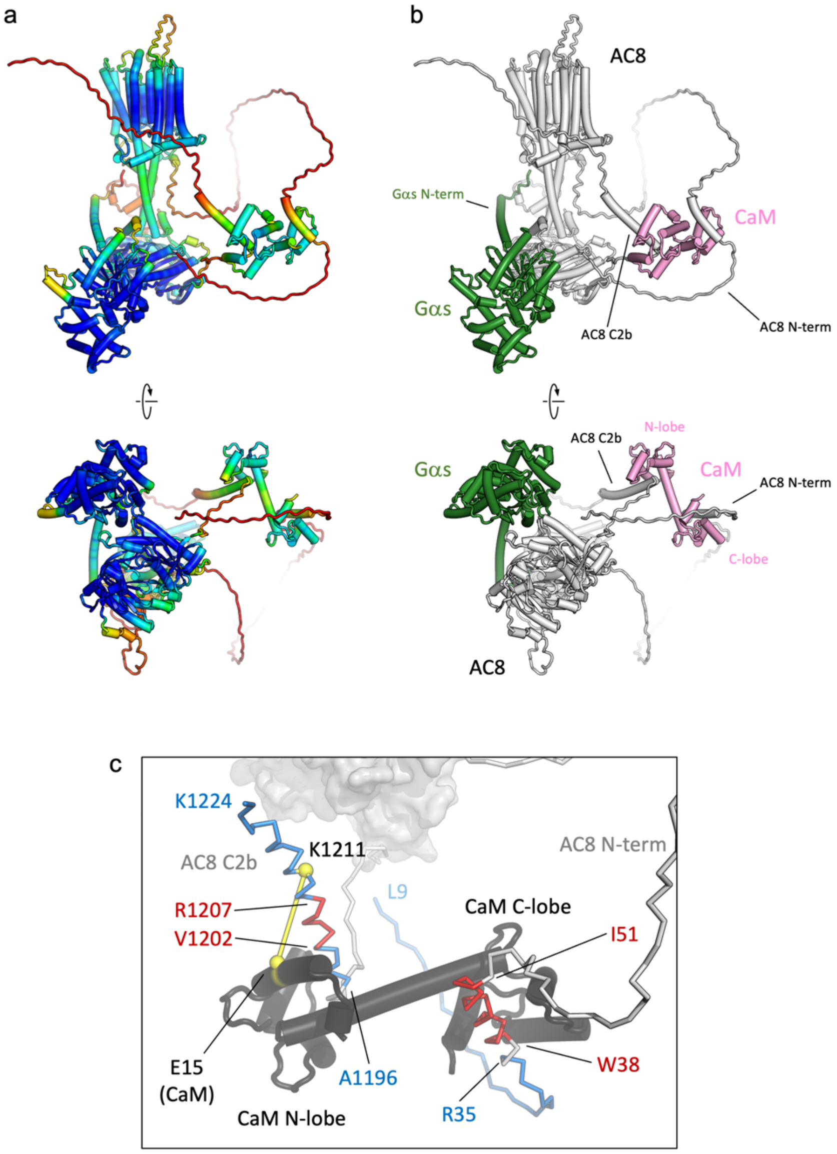


**Figure S16. AlphaFold2 model of AC8-CaM-Gαs. (a)** The views of the model of the complex generated using AlphaFold2 as described in Materials and Methods, coloured according to the pLDDT scores (blue – high pLDDT / confidence; red – low pLDDT / confidence). **(b)** The same views as in a, with key elements of the predicted structure labelled, and individual proteins coloured white (AC8), pink (CaM) and green (Gαs). **(c)** The AlphaFold2 prediction matches well the known CaM binding sites (the 1-5-8-14 motif (WXXXVXXIXXXXXI) residues 38-51; the IQ-like motif (VQXXXR) 1202-1207, red (27)) and the LiP-MS-detected CaM binding peptides (9-35, 1196-1224, blue), respectively, in the N-terminus and C2b domain of AC8. CaM is coloured dark grey. The crosslink between E15 (CaM) and K1211 (AC8) detected in our XL-MS analysis is indicated as a yellow line. The distance between the Cα atoms of these residues (yellow spheres) is 17.5 Å.

**Table S1.** PRM method parameters. Acquisition parameters used for the PRM mass spectrometry method. Higher Orbitrap resolution settings and longer injection times were used for the peptides relevant for absolute quantification.

| Protein | Peptide | Precursor (m/z) | Precursor Charge (z) | Collision Energies (%) | Orbitrap Resolution | Normalized AGC Target (%) | Maximum Injection Time (ms) | Polarity |
|---|---|---|---|---|---|---|---|---|
| (iRT) | ADVTPADFSEWSK (light) | 726.8357 | 2 | 27 | 15000 | 400 | 22 | Positive |
| (iRT) | VEATFGVDESNAK (light) | 683.8279 | 2 | 27 | 15000 | 400 | 22 | Positive |
| (iRT) | YILAGVENSK (light) | 547.298 | 2 | 27 | 15000 | 400 | 22 | Positive |
| (iRT) | DGLDAASYYAPVR (light) | 699.3384 | 2 | 27 | 15000 | 400 | 22 | Positive |
| (iRT) | GAGSSEPVTGLDAK (light) | 644.8226 | 2 | 27 | 15000 | 400 | 22 | Positive |
| (iRT) | GTFIIDPAAVIR (light) | 636.8692 | 2 | 27 | 15000 | 400 | 22 | Positive |
| (iRT) | GTFIIDPGGVIR (light) | 622.8535 | 2 | 27 | 15000 | 400 | 22 | Positive |
| (iRT) | LFLQFGAQGSPFLK (light) | 776.9298 | 2 | 27 | 15000 | 400 | 22 | Positive |
| (iRT) | LGGNEQVTR (light) | 487.2567 | 2 | 27 | 15000 | 400 | 22 | Positive |
| (iRT) | TPVISGGPYEYR (light) | 669.8381 | 2 | 27 | 15000 | 400 | 22 | Positive |
| (iRT) | TPVITGAPYEYR (light) | 683.8537 | 2 | 27 | 15000 | 400 | 22 | Positive |
| CaM | EAFSLFDK (light) | 478.7398 | 2 | 27 | 60000 | 400 | 118 | Positive |
| CaM | EAFSLFDK (heavy) | 482.7469 | 2 | 27 | 60000 | 400 | 118 | Positive |
| AC8 | LLWQTAVR (light) | 493.7927 | 2 | 27 | 60000 | 400 | 118 | Positive |
| AC8 | LLWQTAVR (heavy) | 498.7969 | 2 | 27 | 60000 | 400 | 118 | Positive |
| AC8 | HNIETYLIK (light) | 565.8139 | 2 | 27 | 60000 | 400 | 118 | Positive |
| AC8 | HNIETYLIK (heavy) | 569.821 | 2 | 27 | 60000 | 400 | 118 | Positive |
| AC8 | NILPSHVAR (light) | 503.7933 | 2 | 27 | 60000 | 400 | 118 | Positive |
| AC8 | NILPSHVAR (heavy) | 508.7974 | 2 | 27 | 60000 | 400 | 118 | Positive |
| AC8 | QLLNENNNTGIIK (light) | 735.8992 | 2 | 27 | 60000 | 400 | 118 | Positive |
| AC8 | QLLNENNNTGIIK (heavy) | 739.9063 | 2 | 27 | 60000 | 400 | 118 | Positive |
| CaM | DTDSEEEIR (light) | 547.2358 | 2 | 27 | 60000 | 400 | 118 | Positive |
| CaM | DTDSEEEIR (heavy) | 552.24 | 2 | 27 | 60000 | 400 | 118 | Positive |
| CaM | DGNGYISAAELR (light) | 633.3097 | 2 | 27 | 60000 | 400 | 118 | Positive |
| CaM | DGNGYISAAELR (heavy) | 638.3138 | 2 | 27 | 60000 | 400 | 118 | Positive |

**Table S2.** Cryo-EM analysis and statistics

| **Data collection** | | | | | | |
|---|---|---|---|---|---|---|
| | **AC8-CaM-Gαs In GDN micelle** | | | **AC8-CaM-Gαs In lipid nanodisc** | | |
| | Full | sAC8-Gαs | tmAC8 | Full | sAC8-Gαs | tmAC8 |
| Instrument | FEI Titan Krios / Gatan K3 Summit | | | FEI Titan Krios / Gatan K3 Summit | | |
| Magnification | 130000 | | | 130000 | | |
| Voltage (kV) | 300 | | | 300 | | |
| **Electron Dose (e-/Å2)** | | | | | | |
| Data-set 1 | 60 e-/Å$^2$ | | | 55 e-/Å$^2$ | | |
| Data-set 2 | 56 e-/Å$^2$ | | | 55 e-/Å$^2$ | | |
| Data-set 3 | 49 e-/Å$^2$ | | | 56.4 e-/Å$^2$ | | |
| Defocus range (μm) | -0.6 to -3.0 | | | -0.6 to -3.0 | | |
| Pixel size (Å) | 0.66 | | | 0.66 | | |
| **Refinement** | | | | | | |
| Number of particles | 77575 | | | 150262 | | |
| Map symmetry | C1 | | | C1 | | |
| Model resolution at FSC threshold 0.143 | 3.50 Å | 3.38 Å | 4.13 Å | 3.97 Å | 3.54 Å | 4.20 Å |
| Map sharpening B-factor (Å) | -80 | -80 | -120 | -80 | -80 | -188 |
| Map CC | 0.76 | - | - | | 0.65 | - |
| **Model composition** | | | | | | |
| Protein residues/ligands | 1223/3 | | | 767/3 | | |
| Bond length (r.m.s.d) | 0.010 | | | 0.012 | | |
| Bond angle (r.m.s.d) | 0.630 | | | 0.678 | | |
| **Validation** | | | | | | |
| MolProbity score | 1.92 | | | 2.14 | | |
| Clashscore | 11.73 | | | 17.89 | | |
| Rotamer outlier (%) | 0.46 | | | 0.59 | | |
| **Ramachandran plot** | | | | | | |
| Favored (%) | 95.21 | | | 94.27 | | |
| Allowed (%) | 4.79 | | | 5.73 | | |
| Disallowed (%) | 0 | | | 0 | | |

**Table S3.** Conserved and variable residues in AC-Gαs binding interfaces

| Protein | Gαs | AC8 | AC9 | AC2(C1a)-AC5(c2a) |
|---|---|---|---|---|
| Interactive residues | K223 | - | E1073 | - |
| | Q236 | Q1005 | - | - |
| | Q236 | - | - | N905 |
| | Q236 | - | - | C911 |
| | **N239*** | **R1011** | **R1084** | **R913** |
| | N239 | **-** | **E1081** | **-** |
| | **R280** | **D1019** | **D1092** | **D921** |
| | R280 | E1015 | - | E917 |
| | T284 | **-** | **-** | M378 |

* The conserved interacting residue pairs in all three AC isoforms are bold

### References


1. C. Qi, S. Sorrentino, O. Medalia, V. M. Korkhov, The structure of a membrane adenylyl cyclase bound to an activated stimulatory G protein. *Science* **364**, 389-394 (2019).
2. C. Qi *et al.*, Structural basis of adenylyl cyclase 9 activation. *Nature Communications* **13**, 1045 (2022).
3. R. B. Hitchman, R. D. Possee, L. A. King, in *Recombinant Gene Expression,* A. Lorence, Ed. (Humana Press, Totowa, NJ, 2012), pp. 609-627.
4. M. H. Kubala, O. Kovtun, K. Alexandrov, B. M. Collins, Structural and thermodynamic analysis of the GFP:GFP-nanobody complex. *Protein Science* **19**, 2389-2401 (2010).
5. T. K. Ritchie *et al.*, in *Methods in Enzymology,* N. Düzgünes, Ed. (Academic Press, 2009), vol. 464, pp. 211-231.
6. R. Alvarez, D. V. Daniels, A single column method for the assay of adenylate cyclase. *Analytical Biochemistry* **187**, 98-103 (1990).
7. J. Zivanov *et al.*, New tools for automated high-resolution cryo-EM structure determination in RELION-3. *eLife* **7**, e42166 (2018).
8. S. Q. Zheng *et al.*, MotionCor2: anisotropic correction of beam-induced motion for improved cryo-electron microscopy. *Nature Methods* **14**, 331-332 (2017).
9. K. Zhang, Gctf: Real-time CTF determination and correction. *Journal of Structural Biology* **193**, 1-12 (2016).
10. A. Kucukelbir, F. J. Sigworth, H. D. Tagare, Quantifying the local resolution of cryo-EM density maps. *Nature Methods* **11**, 63-65 (2014).
11. D. Kimanius, L. Dong, G. Sharov, T. Nakane, S. H. W. Scheres, New tools for automated cryo-EM single-particle analysis in RELION-4.0. *Biochemical Journal* **478**, 4169-4185 (2021).
12. R. Bruderer *et al.*, Extending the limits of quantitative proteome profiling with data-independent acquisition and application to acetaminophen-treated three-dimensional liver microtissues. *Mol Cell Proteomics* **14**, 1400-1410 (2015).
13. J. P. Quast, D. Schuster, P. P., protti: an R package for comprehensive data analysis of peptide- and protein-centric bottom-up proteomics data. *Bioinformatics Advances* **2**, (2022).
14. H. Wickham, Welcome to the tidyverse. *Journal of Open Source Software* **4**, 1686 (2019).
15. M. Dowle, A. Srinivasan, data.table: Extension of `data.frame`. https://r-datatable.com, (2022).
16. C. Ritz, F. Baty, J. C. Streibig, D. Gerhard, Dose-Response Analysis Using R. *PLoS One* **10**, e0146021 (2015).
17. E. D. Merkley *et al.*, Distance restraints from crosslinking mass spectrometry: mining a molecular dynamics simulation database to evaluate lysine-lysine distances. *Protein Sci* **23**, 747-759 (2014).
18. A. Leitner *et al.*, Chemical cross-linking/mass spectrometry targeting acidic residues in proteins and protein complexes. *Proc Natl Acad Sci U S A* **111**, 9455-9460 (2014).
19. J. Cox, M. Mann, MaxQuant enables high peptide identification rates, individualized p.p.b.-range mass accuracies and proteome-wide protein quantification. *Nat Biotechnol* **26**, 1367-1372 (2008).
20. A. Leitner, T. Walzthoeni, R. Aebersold, Lysine-specific chemical cross-linking of protein complexes and identification of cross-linking sites using LC-MS/MS and the xQuest/xProphet software pipeline. *Nat Protoc* **9**, 120-137 (2014).
21. B. Schiffrin, S. E. Radford, D. J. Brockwell, A. N. Calabrese, PyXlinkViewer: A flexible tool for visualization of protein chemical crosslinking data within the PyMOL molecular graphics system. *Protein Sci* **29**, 1851-1857 (2020).
22. C. W. Combe, L. Fischer, J. Rappsilber, xiNET: cross-link network maps with residue resolution. *Mol Cell Proteomics* **14**, 1137-1147 (2015).

23. T. L. Davis, T. M. Bonacci, S. R. Sprang, A. V. Smrcka, Structural and molecular characterization of a preferred protein interaction surface on G protein beta gamma subunits. *Biochemistry* **44**, 10593-10604 (2005).
24. G. C. van Zundert, A. M. Bonvin, Modeling protein-protein complexes using the HADDOCK webserver "modeling protein complexes with HADDOCK". *Methods Mol Biol* **1137**, 163-179 (2014).
25. G. C. van Zundert, A. M. Bonvin, DisVis: quantifying and visualizing accessible interaction space of distance-restrained biomolecular complexes. *Bioinformatics* **31**, 3222-3224 (2015).
26. R. Fraczkiewicz, W. Braun, Exact and efficient analytical calculation of the accessible surface areas and their gradients for macromolecules. *Journal of Computational Chemistry* **19**, 319-333 (1998).
27. D. A. MacDougall, S. Wachten, A. Ciruela, A. Sinz, D. M. F. Cooper, Separate Elements within a Single IQ-like Motif in Adenylyl Cyclase Type 8 Impart Ca2+ Calmodulin Binding and Autoinhibition *Journal of Biological Chemistry* **284**, 15573-15588 (2009).
